# Shared network between social exclusion and physiological needs

**DOI:** 10.64898/2026.02.27.707574

**Authors:** J. Bosulu, Y. Mzireg, Y. Luo, S. Hétu

**Affiliations:** Faculté des arts et des sciences, Université de Montréal; School of Psychology and Cognitive Science, East China Normal University

**Keywords:** need, deprivation, meta-analysis, fmri, hunger, thirst, social exclusion, interoception, serotonin, neuroscience

## Abstract

We investigated the neural substrates underlying the brain network shared by social exclusion and physiological needs, both viewed as instances of deprivation. Using activation likelihood estimation (ALE) meta-analyses, we examined brain activation patterns from studies where participants perceived food/water while hungry/thirsty and social interactions while experiencing exclusion. This analysis revealed overlapping activation in the mid-posterior insula, caudate head and ventral anterior cingulate cortex (ACC), as regions consistently engaged when perceiving relevant stimuli across both physiological and social deprivation. Furthermore, we found high spatial correlation between this shared network and the distribution of dopamine receptors, and we identified a significant positive correlation with the 5HT4 among serotonin receptors. Our findings suggest that perceiving deprivation-related stimuli activates brain regions and neurotransmitters involved in aversive affect and goal-directed behavior. These results highlight a neural bridge linking basic physiological drives with complex social needs, offering new insights into the neurobiological architecture of human affective states—while representing just one part of a much larger puzzle.

## INTRODUCTION

Social connection is a fundamental psychological need (Eisenberg, 2012; Reeve & Lee, 2018). Social exclusion represents a form of social disconnection (Eisenberg, 2012) and has been proposed to elicit neural responses similar to those observed during physical pain, particularly through activation of the dorsal anterior cingulate cortex (dACC; Eisenberger, 2012). However, a recent meta-analysis reports consistent activation in the ventral, rather than dorsal, ACC, challenging the interpretation of social exclusion as a pain-like process (Mwilambwe-Tshilobo & Spreng, 2021).

Here, we explore the alternative hypothesis that the brain networks related to social exclusion more closely resemble those associated with unmet physiological needs, such as hunger and thirst, as both can be viewed as instances of deprivation. Indeed, it has been found recently that social isolation shares the same activation of dopaminergic neurons (Tomova et al., 2020). Furthermore, deprivation of biologically significant stimuli—whether physiological (e.g., food or water) or social (e.g., social inclusion)—has long been argued to generate aversive motivational states that guide behavior toward need fulfillment (Maslow, 1943; Hull, 1943; MacGregor, 1960; Baumeister & Leary, 1995; Goebel & Brown, 1981; Hofstede, 1984; Cacioppo et al., 2000; Tay & Diener, 2011; Maner et al., 2007). Neurobiologically, such aversive states are consistently associated with activity in the insula and the anterior cingulate cortex (ACC); whereas the motivational and goal directed aspects are associated with the ventral and dorsal striatum.

Indeed, neuroimaging studies examining responses to stimuli that one is deprived of, i.e. food, water, or social stimuli during states of hunger, thirst, or social exclusion, reveal convergent activation in both the insula (hunger/food: van der Laan et al., 2011; Goldstone et al., 2009; Siep et al., 2009; thirst/water: De Araujo et al., 2003; Egan et al., 2003; Becker et al., 2017; Farrell et al., 2011; social exclusion/social interaction: Masten et al., 2009; Bolling et al., 2011a; Mwilambwe-Tshilobo & Spreng, 2021), the ACC (hunger/food: Goldstone et al., 2009; Siep et al., 2009; Führer et al., 2008; thirst/water: De Araujo et al., 2003; Becker et al., 2015, 2017; Farrell et al., 2011; Saker et al., 2014; social exclusion/social interaction: Eisenberger et al., 2003; Masten et al., 2009; Bolling et al., 2011a; Vijayakumar et al., 2017; Mwilambwe-Tshilobo & Spreng, 2021) and the ventral or dorsal striatum (hunger/food: Bosulu et al., 2022; van der Laan et al., 2011; Siep et al., 2009; social exclusion/social interaction: Masten et al., 2009; Vijayakumar et al., 2017. Despite this overlap, no large scale meta-analysis has quantitatively examined neural substrates of physiological needs and social exclusion that are convergent across studies to determine whether these regions are reliably recruited across both domains.

Furthermore, dopamine (DA) and serotonin (5-HT) interact to mediate behavioral adjustments in response to need states (Boureau & Dayan, 2011). DA is primarily involved in reward prediction, motivation, and action selection (Schultz et al., 1997, 2015; Montague et al., 1996; Schultz, 1998; Rice et al., 2011; Hamid et al., 2016), whereas low 5-HT levels are associated with enhanced processing of aversive, biologically salient events, reflecting neural evaluations of the desirability of the organism’s current state (Dayan & Huys, 2009; Sizemore, 2020; van Galen et al., 2021; Preller et al., 2016; Luo et al., 2016; Liu et al., 2020).

Building on these frameworks, the present study examines whether social exclusion engages neural mechanisms overlapping with those supporting physiological needs. We further investigate the role of DA and 5-HT by assessing how neurotransmitter receptor distributions relate to the neural substrates activated by both social and physiological need states.

## METHODOLOGY

### Meta-Analysis of Brain Coordinate

We employed a meta-analytic approach to quantitatively assess brain activation patterns associated with both physiological (hunger/thirst) and social deprivation (exclusion). Meta-analyses enable the examination of patterns across diverse paradigms, samples, and analytical methods. Deprivation can be conceptualized as comprising two components: the state of deprivation itself (e.g. hunger/thirst/social exclusion) and the specific stimulus that is lacking (e.g. food/water/social interaction) (Bindra, 1974; Toates, 1994). Accordingly, we first conducted two meta-analyses: one summarizing fMRI studies on physiological deprivation (hunger/food and thirst/water), focusing on participants perceiving food while hungry or water while thirsty (hereafter referred to as Physiological Deprivation); and another on social deprivation, examining studies where participants viewed social interactions they were excluded from (referred to as Social Deprivation). We then conducted contrast and conjunction analyses to highlight both distinct and overlapping neural activation patterns between the two forms of deprivation. The last meta-analysis allowed us to identify brain regions consistently involved in processing stimuli related to both physiological and social deprivation.

We followed the PRISMA framework to identify relevant articles. For hunger and thirst we ran two searches with different keywords: for hunger: “hunger” OR “food deprivation” AND (“fMRI”) and for thirst: “thirst” OR “water deprivation” AND (“fMRI”. For social exclusion, we used the keywords (“social” AND “exclusion” AND “fMRI”). Searches were conducted on Google Scholar and PubMed, supplemented by reference lists and review articles.

Inclusion criteria for both physiological and social deprivation studies were: healthy participants, whole-brain analyses (with or without small volume correction (SVC)), MNI or Talairach coordinates (converted to MNI SPM152 using the Lancaster transform; Lancaster et al., 2007), corrected activation maps (or cluster-level corrections), and activation contrasts only (see Table 1).

**Table 1.** Selection Criteria.

| Criteria | Hunger and Thirst | Social exclusion |
| --- | --- | --- |
| Privation contrast | Yes | Yes |
| Presence of the relevant rewarding cue | Yes | Yes |
| Passive viewing | Yes | Yes |
| Long term (in hours) deprivation implicating physiological factors | Yes | No |
| Healthy individuals only | Yes | Yes |
| MNI or Talairach Coordinates | Yes | Yes |
| Whole brain contrast (with or without SVC) | Yes | Yes |
| Corrected | Yes | Yes |
| Activation contrast only | Yes | Yes |
| Excluded: MRI and resting states; cognitive conjunction analysis; and functional connectivity results | Yes | Yes |

Table indicating selection criteria. The red coloured “yes” or “no” means the criterion is crucial for the definition of either ‘physiological or social deprivation.

The search yielded 165 articles for hunger and 19 for thirst, with an additional 7 and 3 articles identified through references, respectively. Importantly, for physiological deprivation studies, we applied two additional criteria: (1) participants were in a food or water-deprived state, and (2) participants were exposed to food (while hungry) or water (while thirsty) stimuli via visual, taste, or olfactory cues. The scan results were compared to perception of food/water under satiety. Ultimately, 17 hunger-related and 4 thirst-related articles met all inclusion criteria (see Tables 2 and 3).

**Table 2.** Hunger Selected Articles.

| Paper | Contrast | Stimuli | Task | Healthy Participants |
| --- | --- | --- | --- | --- |
| Jiang, T., Soussignan, R., Schaal, B., & Royet, J. P. (2014). Reward for food odors: an fMRI study of liking | Liking – Wanting (hunger)<br>Wanting – Liking (hunger) | Odor | Odor presentation and rating | 12 |
| and wanting as a function of metabolic state and BMI. Social cognitive and affective neuroscience, 10(4), 561-568. | Food - NFood (hunger) |  |  |  |
| Green, E., Jacobson, A., Haase, L., & Murphy, C. (2015). Neural correlates of taste and pleasantness evaluation in the metabolic syndrome. Brain research, 1620, 57-71. | Hunger-Satiety: Control Sucrose>Caffeine (during hunger) Caffeine>Sucrose (hunger) | Taste | Swallowing aqueous solution | 15 |
| Martens, M. J., Born, J. M., Lemmens, S. G., Karhunen, L., Heinecke, A., Goebel, R., ... & Westerterp-Plantenga, M. S. (2013). Increased sensitivity to food cues in the fasted state and decreased inhibitory control in the satiated state in the overweight. The American journal of clinical nutrition, 97(3), 471-479. | Fasted: F>NF<br>Fasted: stimuli - subject group<br>Fasted: correlation F>NF with BMI | Visual | Viewing food and non food pictures | 40 |
| LaBar, K. S., Gitelman, D. R., Parrish, T. B., Kim, Y. H., Nobre, A. C., & Mesulam, M. (2001). Hunger selectively modulates corticolimbic activation to food stimuli in humans. Behavioral neuroscience, 115(2), 493. | Hungry-Satiated | Visual | Viewing food and tool images | 17 |
| Harding, I. H., Andrews, Z. B., Mata, F., Orlandea, S., Martinez-Zalacain, I., Soriano-Mas, C., ... & Verdejo-Garcia, A. (2018). | Healthy versus unhealthy food choice: fasted>satiated | Visual | participants were asked to select an option using a two-button response box. | 30 |
| Brain substrates of unhealthy versus healthy food choices: influence of homeostatic status and body mass index. International Journal of Obesity, 42(3), 448. |  |  |  |  |
| Frank, S., Laharnar, N., Kullmann, S., Veit, R., Canova, C., Hegner, Y. L., ... & Preissl, H. (2010). Processing of food pictures: influence of hunger, gender and calorie content. Brain research, 1350, 159-166. | HiCal hungry vs. HiCal satiated | Visual | One-back task: press a button, either to indicate that the seen image was the same or another button to indicate that the picture was not the same | 12 |
| Holsen, L. M., Zarcone, J. R., Thompson, T. I., Brooks, W. M., Anderson, M. F., Ahluwalia, J. S., ... & Savage, C. R. (2005). Neural mechanisms underlying food motivation in children and adolescents. Neuroimage, 27(3), 669-676. | Premeal : Food > non-food | Visual | Viewing pictures of food, animals, and baseline control images | 9 |
| Jacobson, A., Green, E., Haase, L., Szajer, J., & Murphy, C. (2019). Differential Effects of BMI on Brain Response to Odor in Olfactory, Reward and Memory Regions: Evidence from fMRI. Nutrients, 11(4), 926. | Odor during the hunger condition | Odor/Taste | Odor stimuli delivered to the tongue | 40 |
| Haase, L., Green, E., & Murphy, C. (2011). Males and females show differential brain activation to taste when hungry and sated in gustatory and reward areas. *Appetite*, 57(2), 421-434. | Hunger x male x Sucrose > water<br>Hunger x female x NaCl > water<br>Hunger x female x CAFFEINE > water<br>Hunger x female x SUCROSE > water<br>Hunger x female x CITRIC ACID > water | Taste | Taste stimuli presentation | 21 |
| Führer, D., Zysset, S., & Stumvoll, M. (2008). Brain activity in hunger and satiety: an exploratory visually stimulated FMRI study. *Obesity*, 16(5), 945-950. | Hunger > Satiety | Visual | Viewing pictures and two-back task: the subject was asked to press a button when the same letter was shown two steps earlier | 12 |
| Haase, L., Cerf-Ducastel, B., & Murphy, C. (2009). Cortical activation in response to pure taste stimuli during the physiological states of hunger and satiety. *Neuroimage*, 44(3), 1008-1021. | Hunger > satiety x sucrose<br>Hunger > satiety x Caffeine<br>Hunger > satiety x Saccharin<br>Hunger > satiety x Citric acid | Taste | Stimulus presentation delivered to the tip of the tongue | 18 |
| Uher, R., Treasure, J., Heining, M., Brammer, M. J., & Campbell, I. C. (2006). Cerebral processing of food-related stimuli: effects of fasting and gender. *Behavioural brain research*, 169(1), 111-119. | Fasting > satiety x food-related stimuli (Chocolate AND Chicken)<br>Fasting > satiety x food-related stimuli (Chocolate)<br>Fasting > satiety x food-related stimuli | Visual | Viewing photographs of food | 18 |
|  | (Chicken) |  |  |  |
| Holsen, L. M., Zarcone, J. R., Brooks, W. M., Butler, M. G., Thompson, T. I., Ahluwalia, J. S., ... & Savage, C. R. (2006). Neural mechanisms underlying hyperphagia in Prader-Willi syndrome. Obesity, 14(6), 1028-1037. | HW x Pre-meal x food > non-food<br>Hw x Pre-meal x non-food > food | Visual | Viewing pictures of food, animals and control | 9 |
| Jacobson, A., Green, E., & Murphy, C. (2010). Age-related functional changes in gustatory and reward processing regions: An fMRI study. Neuroimage, 53(2), 602-610. | Hunger x Older adults x Sucrose<br>Hunger x Young adults x Sucrose<br>Hunger x Older adults x Citric acid<br>Hunger x Younger adults x Citric acid<br>Hunger x Older adults x NaCl<br>Hunger x Younger adults x NaCl<br>Hunger x Older adults x Caffeine<br>Hunger x Younger adults x Caffeine | Taste | Stimuli were delivered orally | 38 |
| Cheah, Y. S., Lee, S., Ashoor, G., Nathan, Y., Reed, L. J., Zelaya, F. O., ... & Amiel, S. A. (2014). Ageing diminishes the modulation of human brain responses to visual food cues by meal ingestion. International Journal of | FASTED > FED x Visual food cue | Visual | Viewing food and non food | 24 |
| Obesity, 38(9), 1186. |  |  |  |  |
| He, Q., Huang, X., Zhang, S., Turel, O., Ma, L., & Bechara, A. (2019). Dynamic causal modeling of insular, striatal, and prefrontal cortex activities during a food-specific Go/NoGo task. *Biological Psychiatry: Cognitive Neuroscience and Neuroimaging*, 4(12), 1080-1089. | Hungry > Satiated | Visual | Food pictures | 45 |
| Kerem, L., Holsen, L., Fazeli, P., Bredella, M. A., Mancuso, C., Resulaj, M., ... & Lawson, E. A. (2021). Modulation of neural fMRI responses to visual food cues by overeating and fasting interventions: A preliminary study | Hungry > Satiated | Visual | Food pictures | 6 |

List of articles that were selected. for hunger..

**Table 3.**
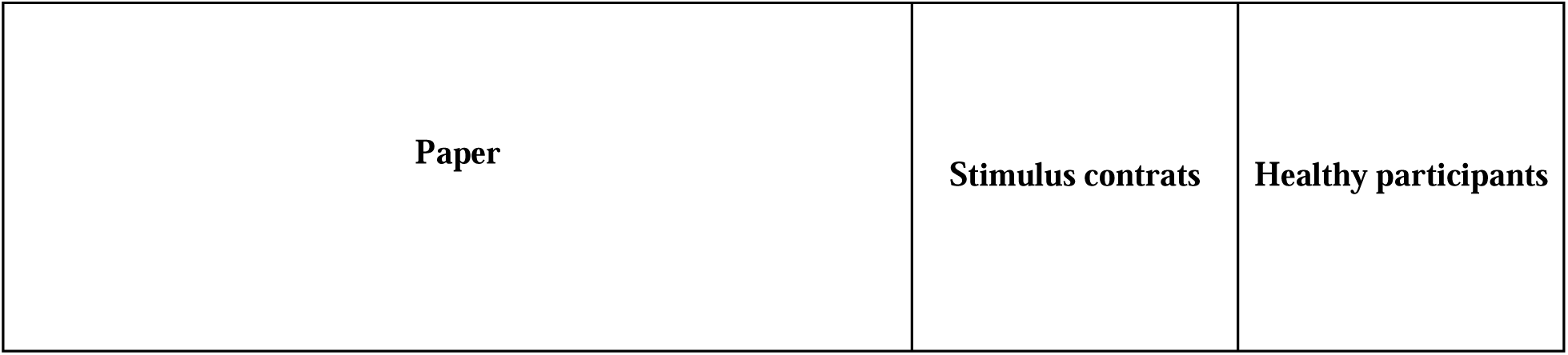
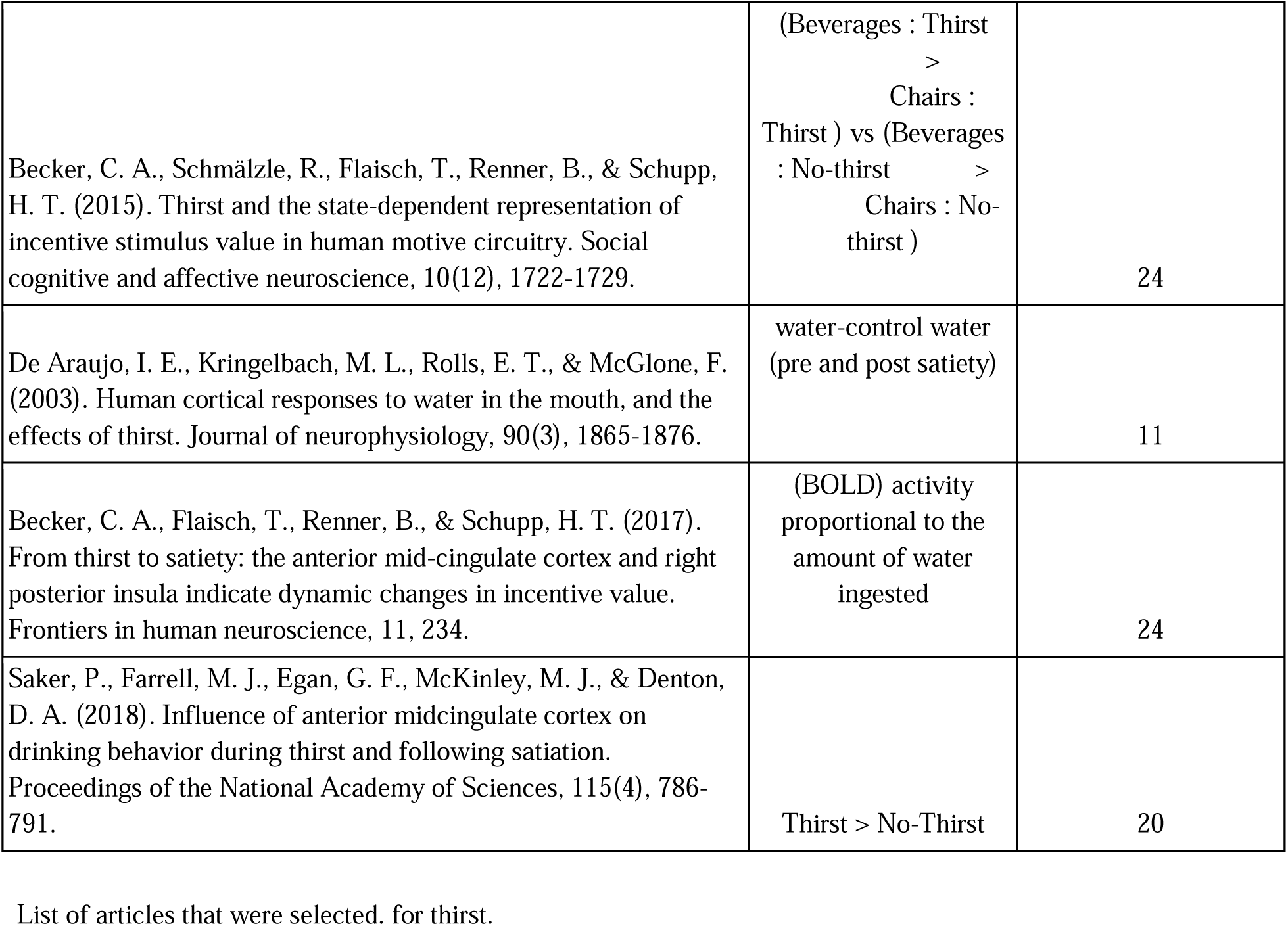
Thirst Selected Articles.

| Paper | Stimulus contrasts | Healthy participants |
| --- | --- | --- |
| Becker, C. A., Schmälzle, R., Flaisch, T., Renner, B., & Schupp, H. T. (2015). Thirst and the state-dependent representation of incentive stimulus value in human motive circuitry. *Social cognitive and affective neuroscience*, 10(12), 1722-1729. | (Beverages : Thirst<br>><br>Chairs :<br>Thirst ) vs (Beverages<br>: No-thirst ><br>Chairs : No-thirst ) | 24 |
| De Araujo, I. E., Kringelbach, M. L., Rolls, E. T., & McGlone, F. (2003). Human cortical responses to water in the mouth, and the effects of thirst. *Journal of neurophysiology*, 90(3), 1865-1876. | water-control water<br>(pre and post satiety) | 11 |
| Becker, C. A., Flaisch, T., Renner, B., & Schupp, H. T. (2017). From thirst to satiety: the anterior mid-cingulate cortex and right posterior insula indicate dynamic changes in incentive value. *Frontiers in human neuroscience*, 11, 234. | (BOLD) activity<br>proportional to the<br>amount of water<br>ingested | 24 |
| Saker, P., Farrell, M. J., Egan, G. F., McKinley, M. J., & Denton, D. A. (2018). Influence of anterior midcingulate cortex on drinking behavior during thirst and following satiation. *Proceedings of the National Academy of Sciences*, 115(4), 786-791. | Thirst > No-Thirst | 20 |

List of articles that were selected. for thirst.

For social deprivation, we identified 130 articles, selecting 22, with 4 additional articles found through reference checks and reviews. Similar to physiological deprivation, two additional criteria were applied: (1) participants were in a state of social deprivation (isolated or excluded), and (2) participants were exposed to social interactions from which they were excluded. The scan results when socially excluded were compared to those where they were included. Notably, all included studies employed the Cyberball task—a virtual ball-tossing game designed to simulate short-term social exclusion (Williams et al., 2000). This paradigm reliably induces emotional distress, reflecting a threat to fundamental social needs (Williams, 2007), and has been proposed to trigger need-like emotional distress (Bernstein & Claypool, 2012).. Applying these criteria, we selected 26 articles on social deprivation for the meta-analysis (see Table 4).

**Table 4.**
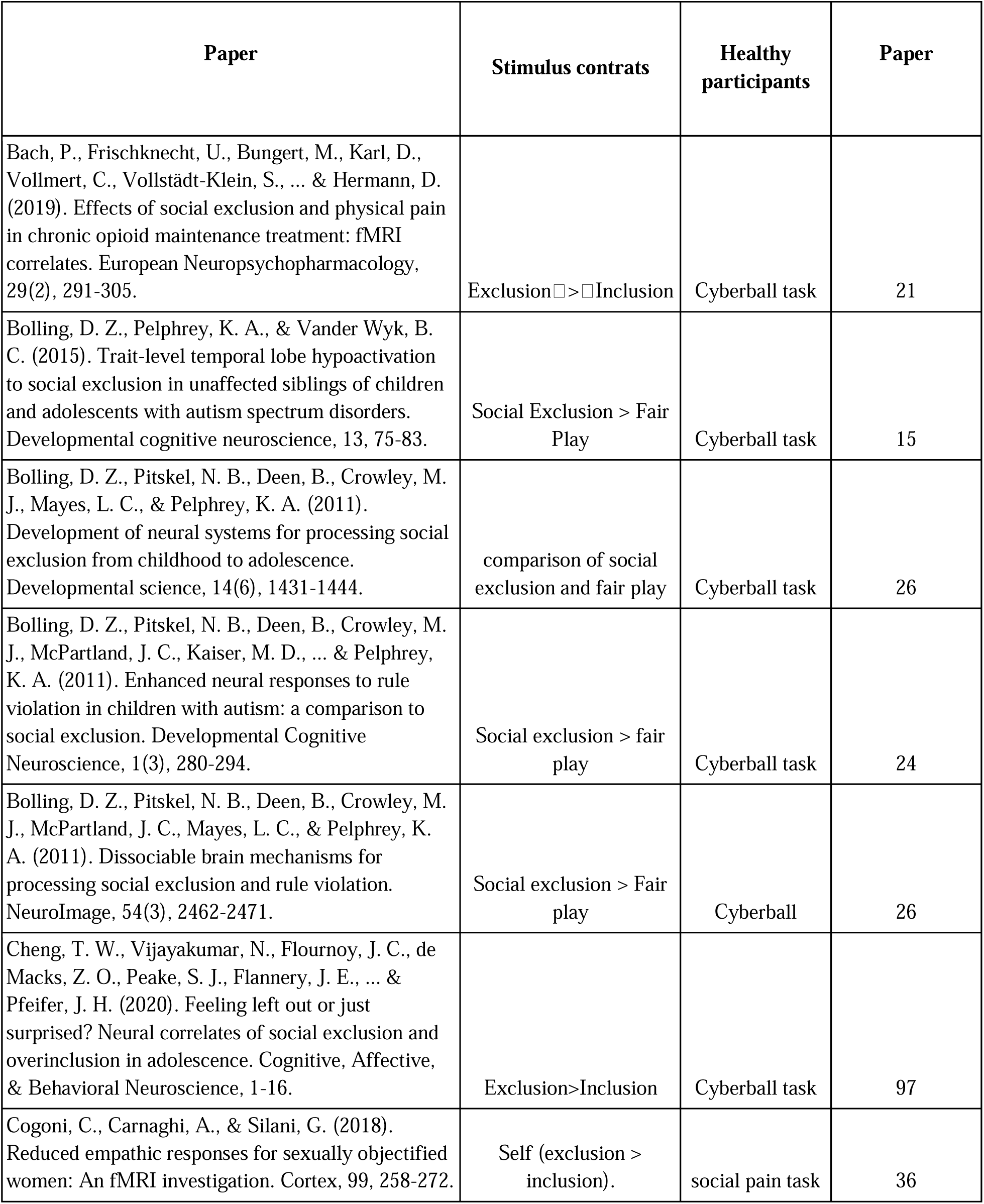
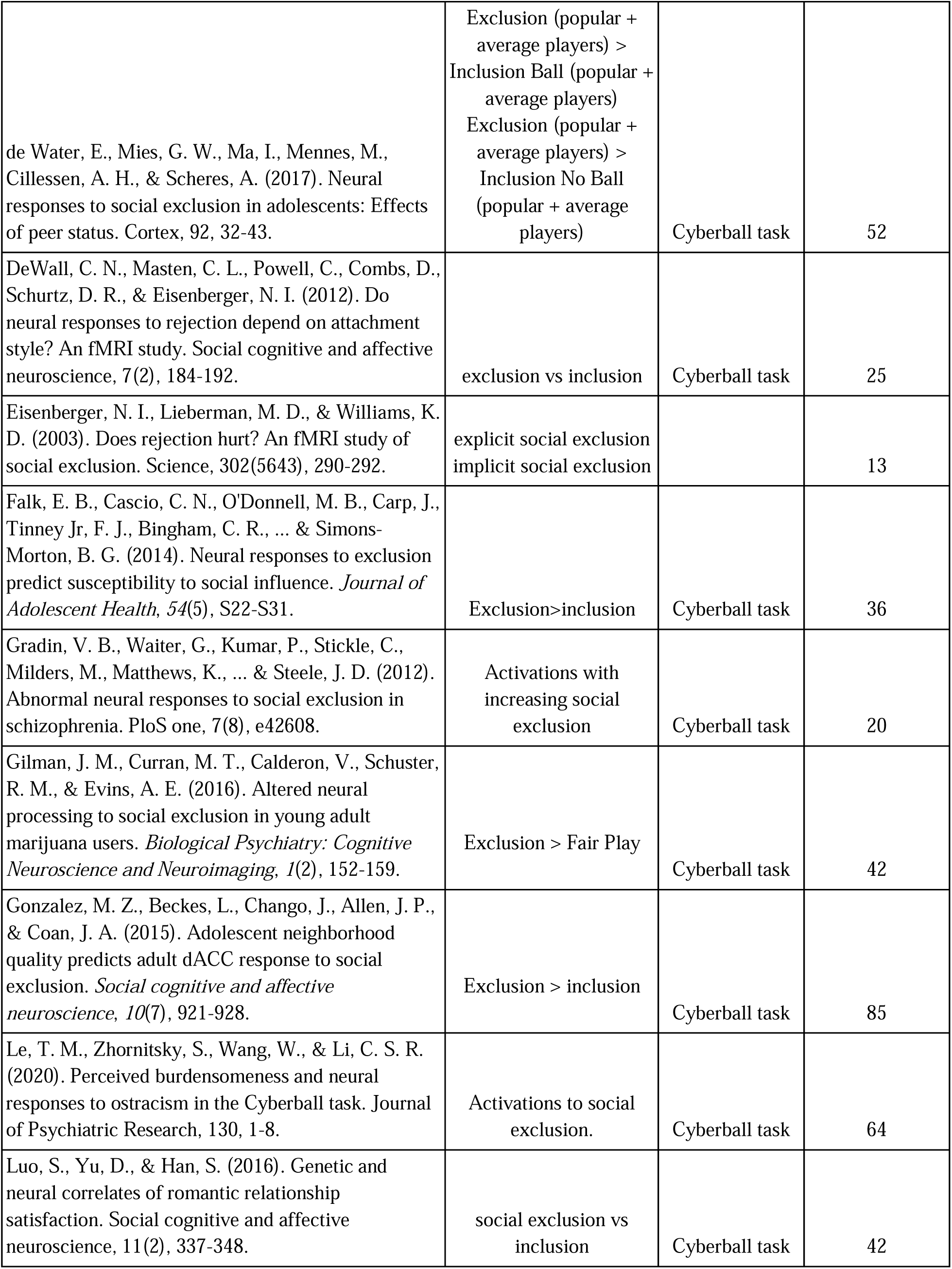
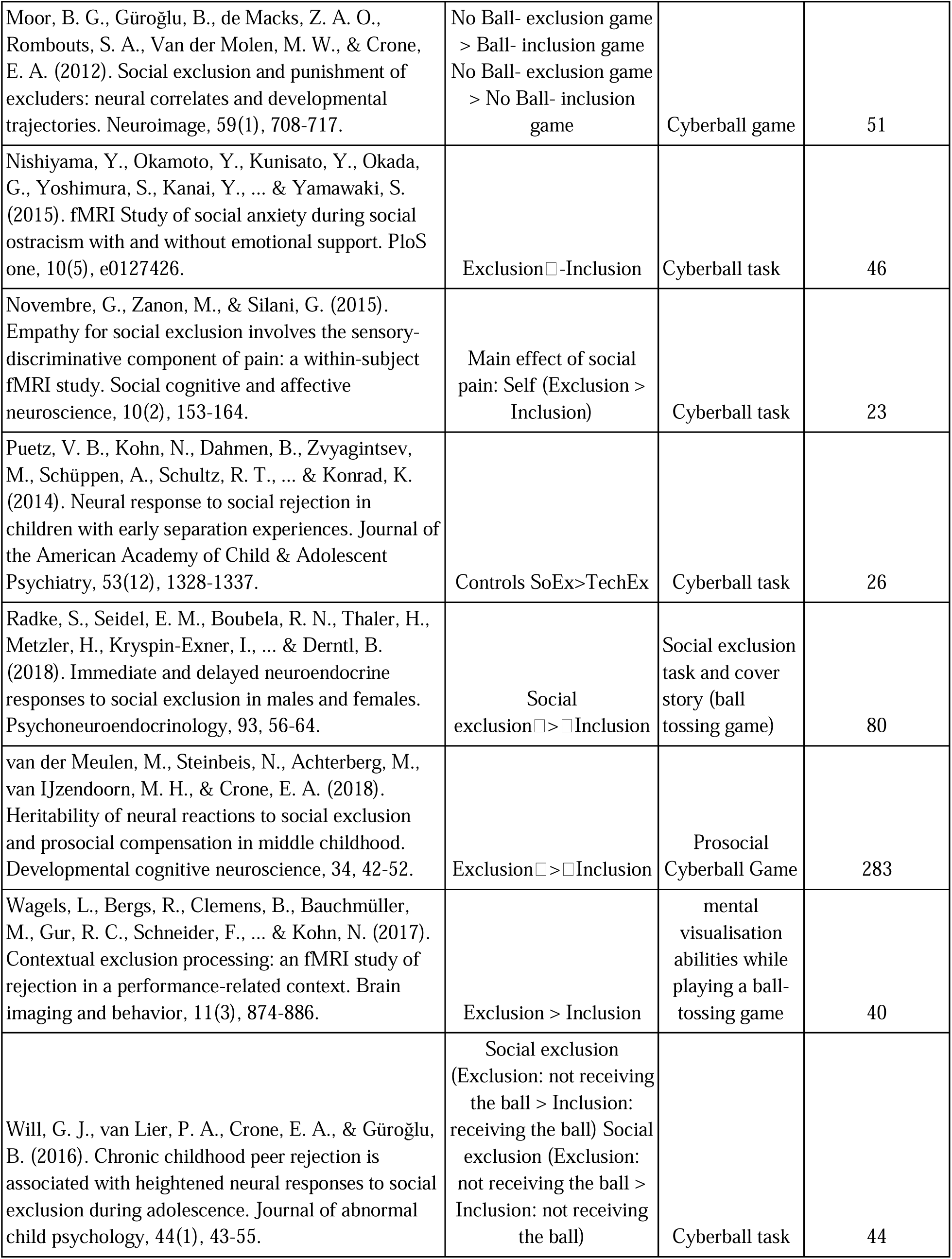
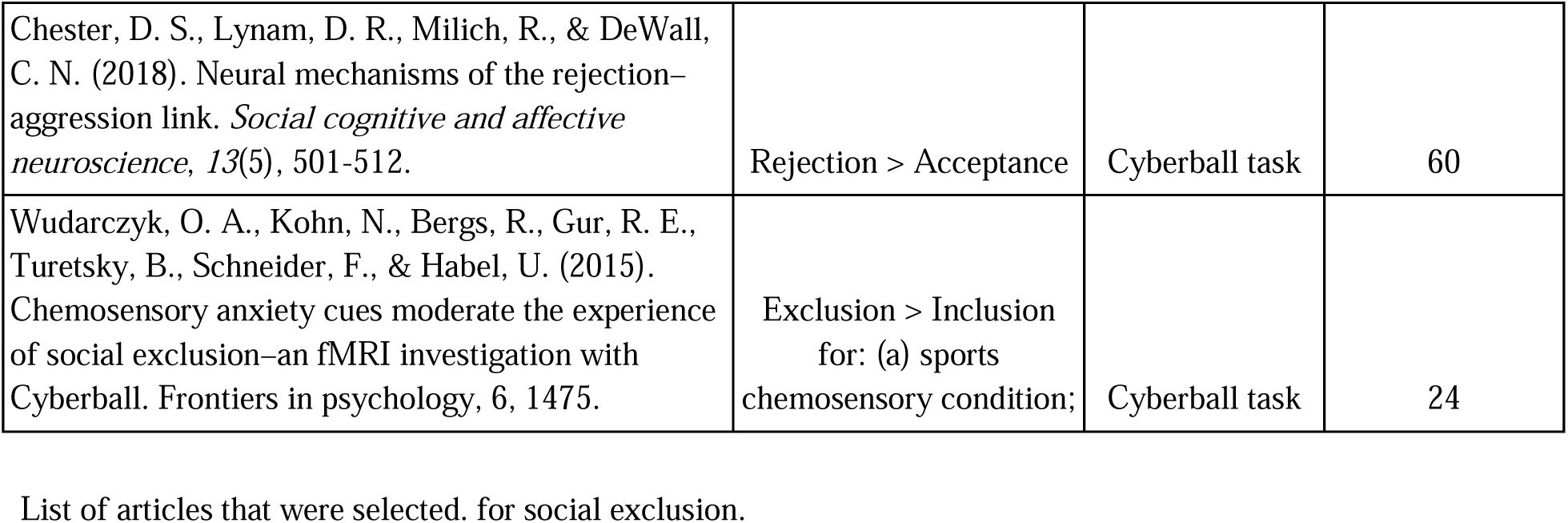
Social Deprivation Selected Articles.

Meta-analyses of functional neuroimaging studies were conducted using the activation likelihood estimation (ALE) approach in GingerALE (Brainmap; Eickhoff et al., 2009, 2012). This method models activation foci as spatial probability distributions, accounting for variability across studies and subjects. ALE maps were generated by computing the union of these probabilities and thresholded using cluster-level FWE correction (P < 0.05) with a voxel-level cluster-forming threshold of P < 0.001 (1000 permutations; Eklund et al., 2016; Woo et al., 2014). Analyses were performed in MNI152 space using an anatomically constrained mask. For physiological deprivation (hunger/thirst, 20 articles, 44 experiments, 856 subjects, 612 foci) and social deprivation (26 articles, 33 experiments, 1511 subjects, 342 foci), we performed contrast and conjunction analyses to identify distinct and shared activations, respectively. All maps were visualized using Mango. (http://ric.uthscsa.edu/mango/).

### Spatial Correlation with Neurotransmitters

We investigated spatial relationships between the intersection of physiological and social need maps and receptor distributions (JuSpace toolbox, Dukart et al., 2021). Using z-transformed correlations (corrected for spatial autocorrelation via SPM12’s TPM.nii), we tested for significant associations between subregions of the intersection map and receptor maps against a null distribution using one-sample t-tests (Dukart et al., 2021).

## RESULTS

### Single Meta-Analyses

The meta-analysis of physiological deprivation, combining hunger and thirst, revealed consistent activation in the bilateral anterior, middle, and posterior insula; bilateral claustrum and putamen, bilateral ACC (Brodmann areas (BA) 24/25), bilateral caudate head and right caudate body, left parahippocampal gyrus, medial frontal gyrus, mammillary body, and hippocampus (Table 5, Figure 1).

**Table 5.**
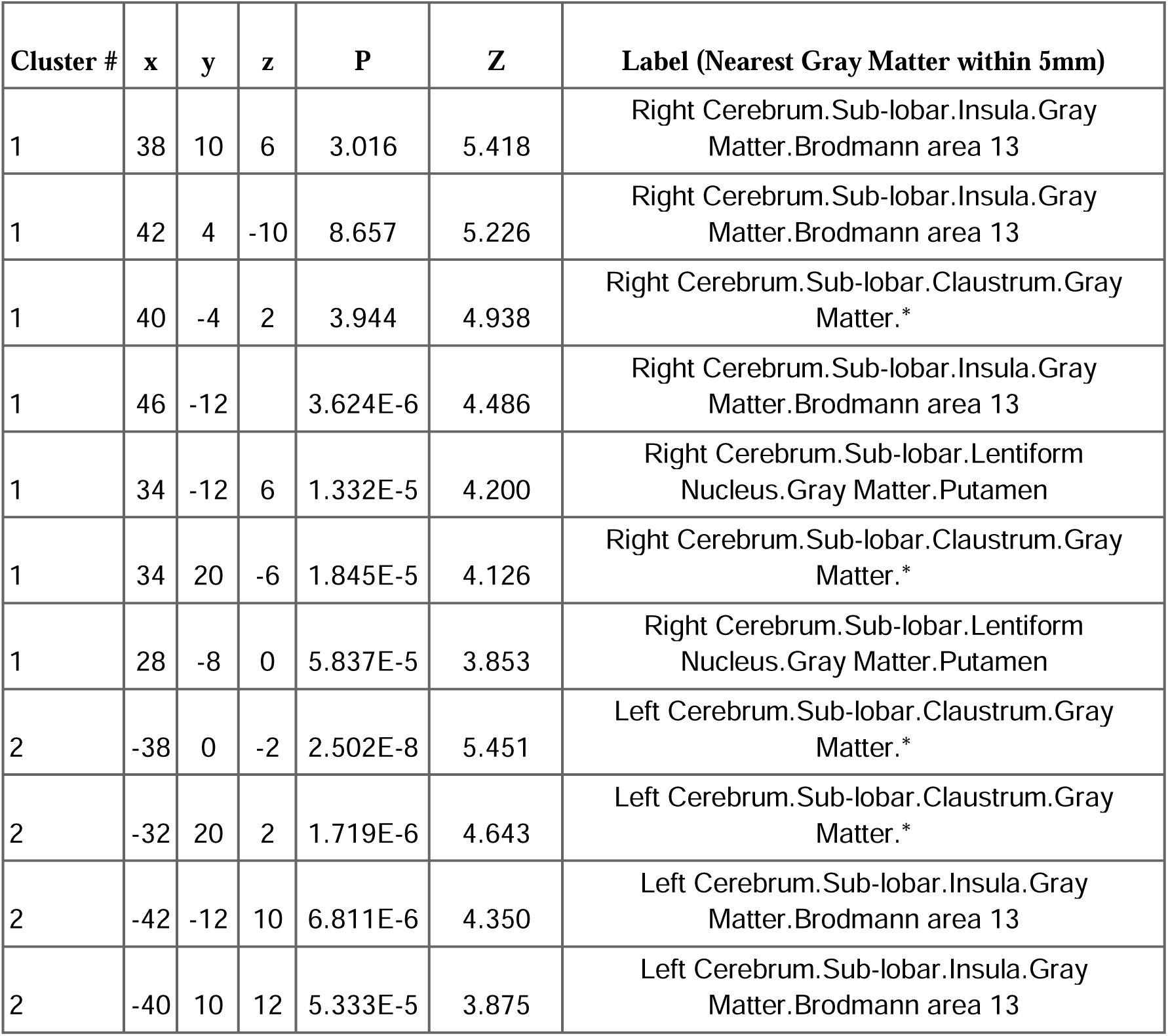
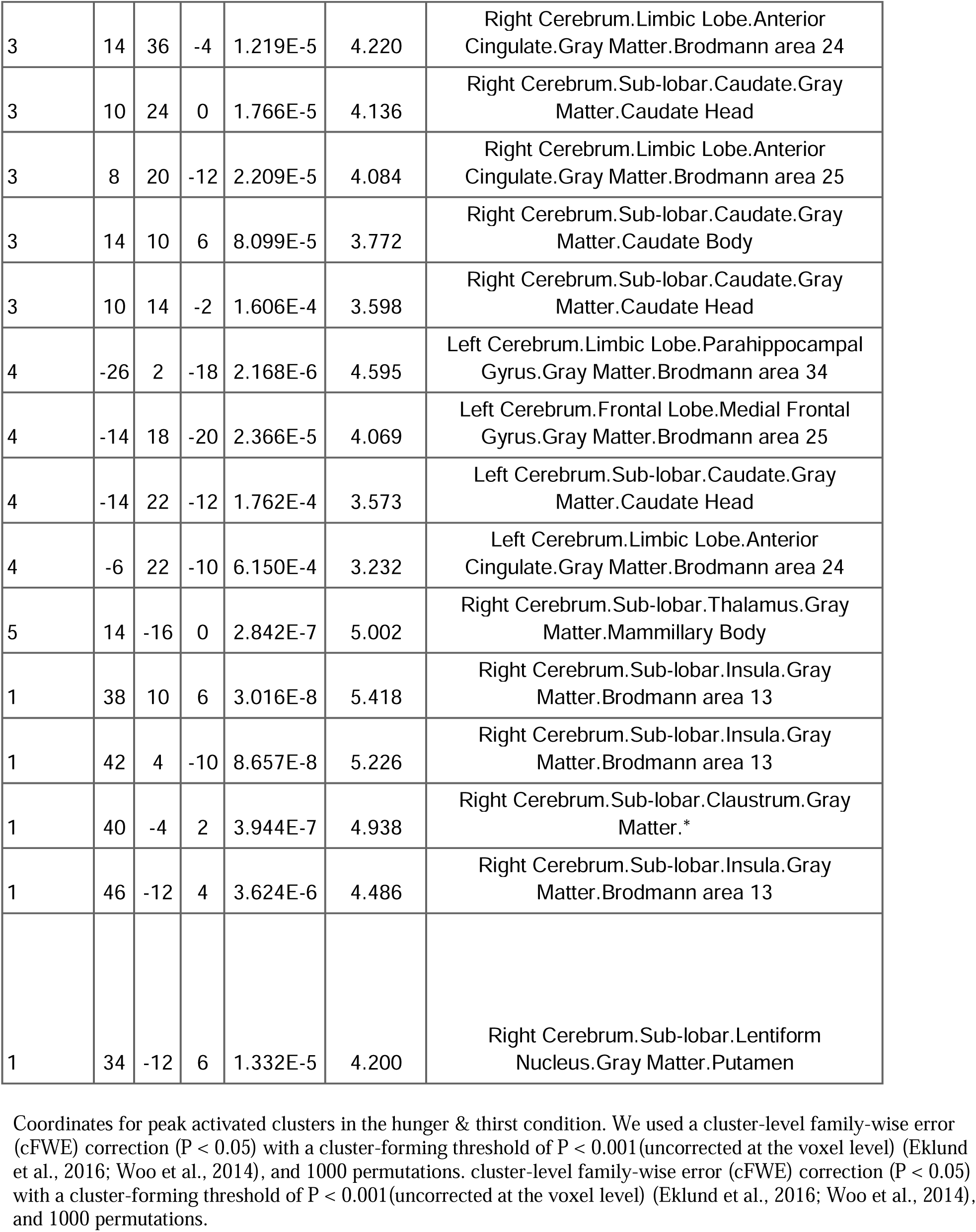
Hunger & Thirst.

**Figure 1.**
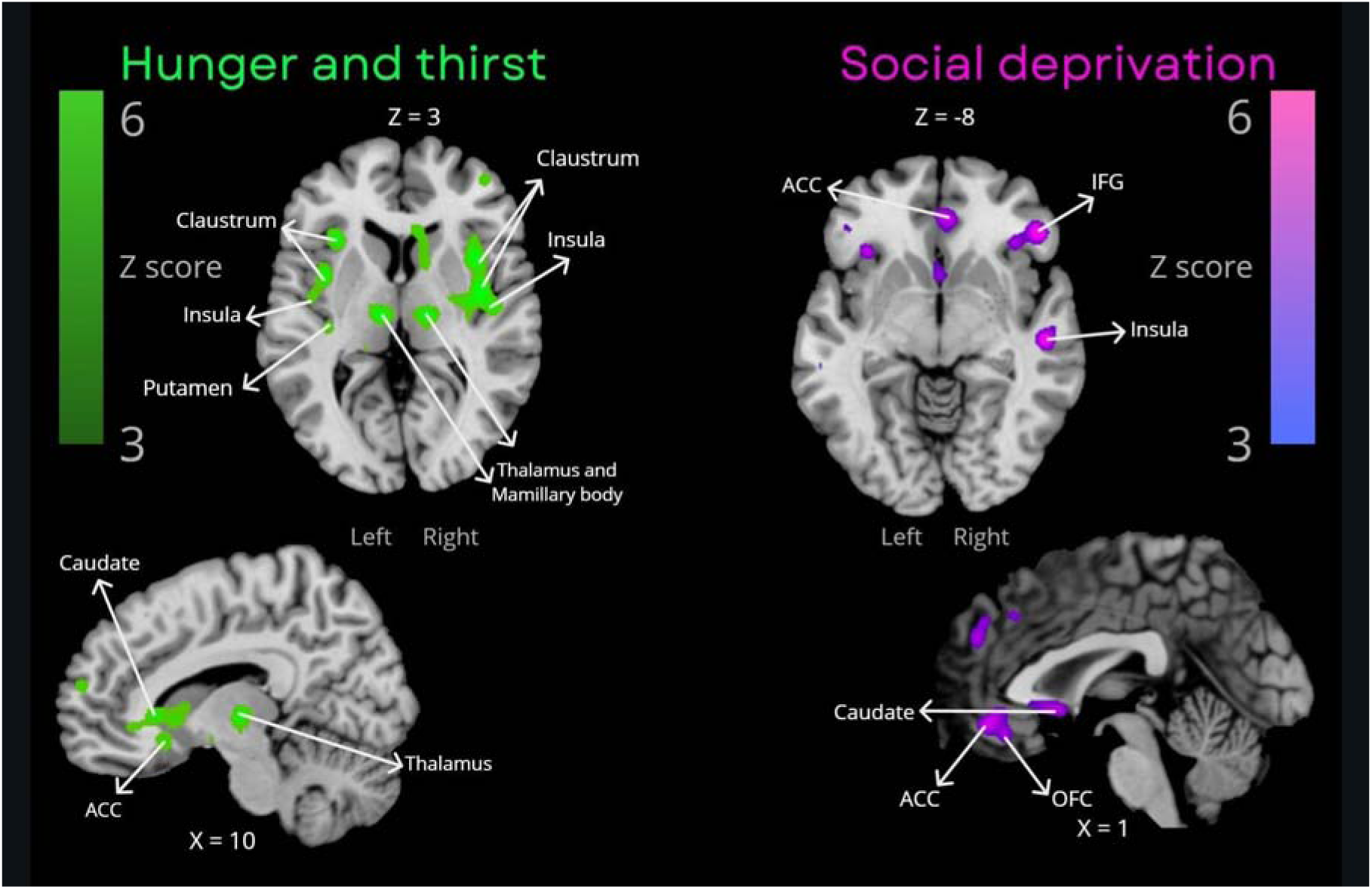
Single meta-analyses maps. Maps for consistently activated clusters in physiological deprivation (green) and social deprivation (pink).

The meta-analysis of social deprivation revealed consistent activation in the right anterior insula; bilateral posterior insula, bilateral ACC (Brodmann areas 24/32), right inferior frontal gyrus, OFC, left anterior transverse temporal gyrus, and bilateral caudate head (Table 6, Figure 1).

**Table 6.**
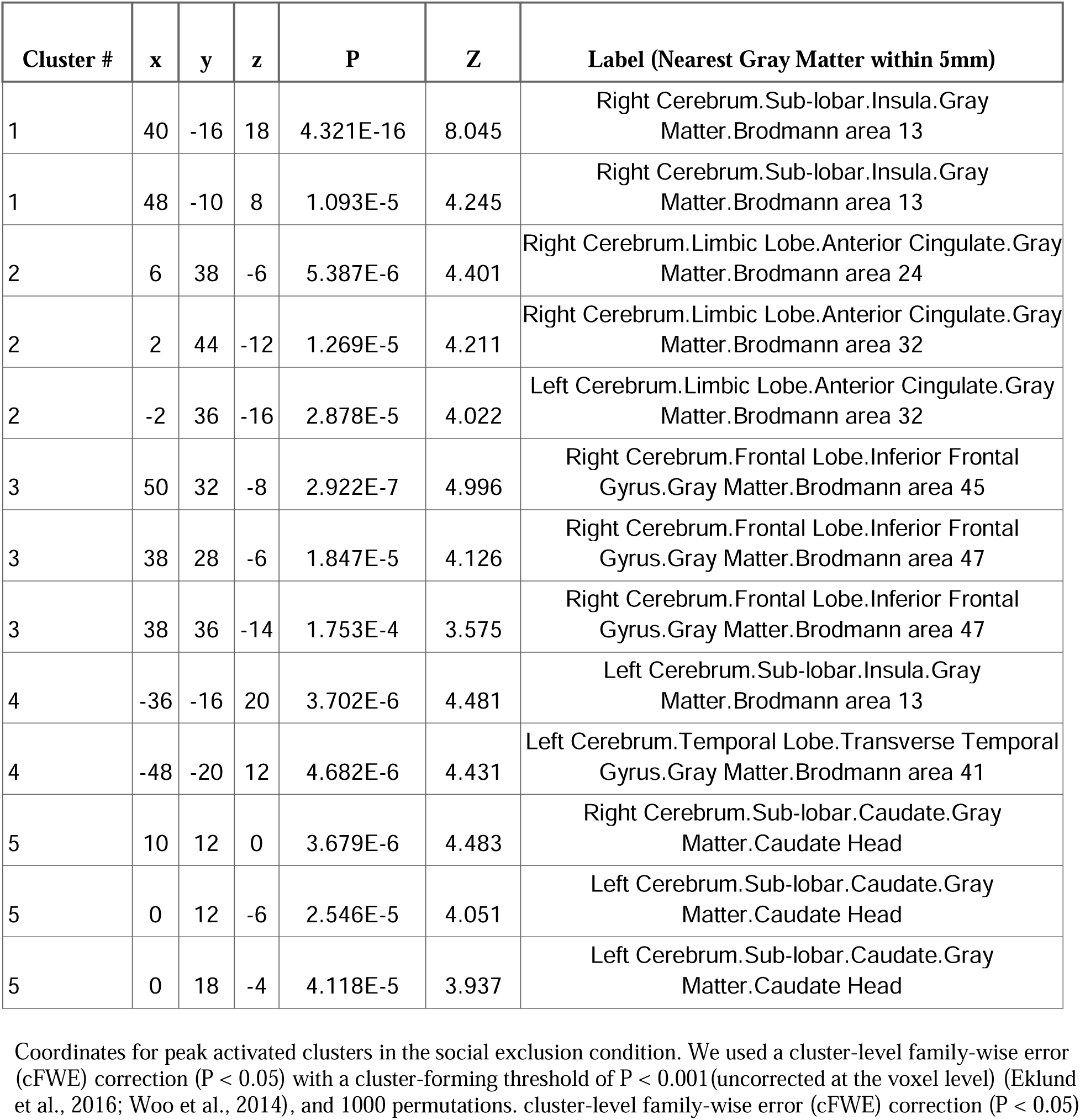
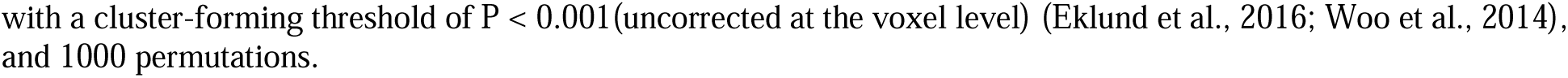
Social Exclusion.

| Cluster # | x | y | z | P | Z | Label (Nearest Gray Matter within 5mm) |
| --- | --- | --- | --- | --- | --- | --- |
| 1 | 40 | -16 | 18 | 4.321E-16 | 8.045 | Right Cerebrum.Sub-lobar.Insula.Gray Matter.Brodmann area 13 |
| 1 | 48 | -10 | 8 | 1.093E-5 | 4.245 | Right Cerebrum.Sub-lobar.Insula.Gray Matter.Brodmann area 13 |
| 2 | 6 | 38 | -6 | 5.387E-6 | 4.401 | Right Cerebrum.Limbic Lobe.Anterior Cingulate.Gray Matter.Brodmann area 24 |
| 2 | 2 | 44 | -12 | 1.269E-5 | 4.211 | Right Cerebrum.Limbic Lobe.Anterior Cingulate.Gray Matter.Brodmann area 32 |
| 2 | -2 | 36 | -16 | 2.878E-5 | 4.022 | Left Cerebrum.Limbic Lobe.Anterior Cingulate.Gray Matter.Brodmann area 32 |
| 3 | 50 | 32 | -8 | 2.922E-7 | 4.996 | Right Cerebrum.Frontal Lobe.Inferior Frontal Gyrus.Gray Matter.Brodmann area 45 |
| 3 | 38 | 28 | -6 | 1.847E-5 | 4.126 | Right Cerebrum.Frontal Lobe.Inferior Frontal Gyrus.Gray Matter.Brodmann area 47 |
| 3 | 38 | 36 | -14 | 1.753E-4 | 3.575 | Right Cerebrum.Frontal Lobe.Inferior Frontal Gyrus.Gray Matter.Brodmann area 47 |
| 4 | -36 | -16 | 20 | 3.702E-6 | 4.481 | Left Cerebrum.Sub-lobar.Insula.Gray Matter.Brodmann area 13 |
| 4 | -48 | -20 | 12 | 4.682E-6 | 4.431 | Left Cerebrum.Temporal Lobe.Transverse Temporal Gyrus.Gray Matter.Brodmann area 41 |
| 5 | 10 | 12 | 0 | 3.679E-6 | 4.483 | Right Cerebrum.Sub-lobar.Caudate.Gray Matter.Caudate Head |
| 5 | 0 | 12 | -6 | 2.546E-5 | 4.051 | Left Cerebrum.Sub-lobar.Caudate.Gray Matter.Caudate Head |
| 5 | 0 | 18 | -4 | 4.118E-5 | 3.937 | Left Cerebrum.Sub-lobar.Caudate.Gray Matter.Caudate Head |

Coordinates for peak activated clusters in the social exclusion condition. We used a cluster-level family-wise error (cFWE) correction ( $P < 0.05$ ) with a cluster-forming threshold of $P < 0.001$ (uncorrected at the voxel level) (Eklund et al., 2016; Woo et al., 2014), and 1000 permutations. cluster-level family-wise error (cFWE) correction ( $P < 0.05$ )
with a cluster-forming threshold of $P < 0.001$ (uncorrected at the voxel level) (Eklund et al., 2016; Woo et al., 2014), and 1000 permutations.

### Contrasts Meta-Analyses

The results of the contrast meta-analyses are presented in Tables 7 and 8 and Figure 2. Compared to perceiving social interactions while socially deprived, perceiving food or water while hungry or thirsty showed more consistent activation in the right putamen, claustrum, bilateral posterior insula, right OFC, and bilateral caudate. Conversely, perceiving social interactions during social deprivation, compared to perceiving food or water during hunger or thirst, elicited more consistent activation in the right posterior insula.

**Table 7.**
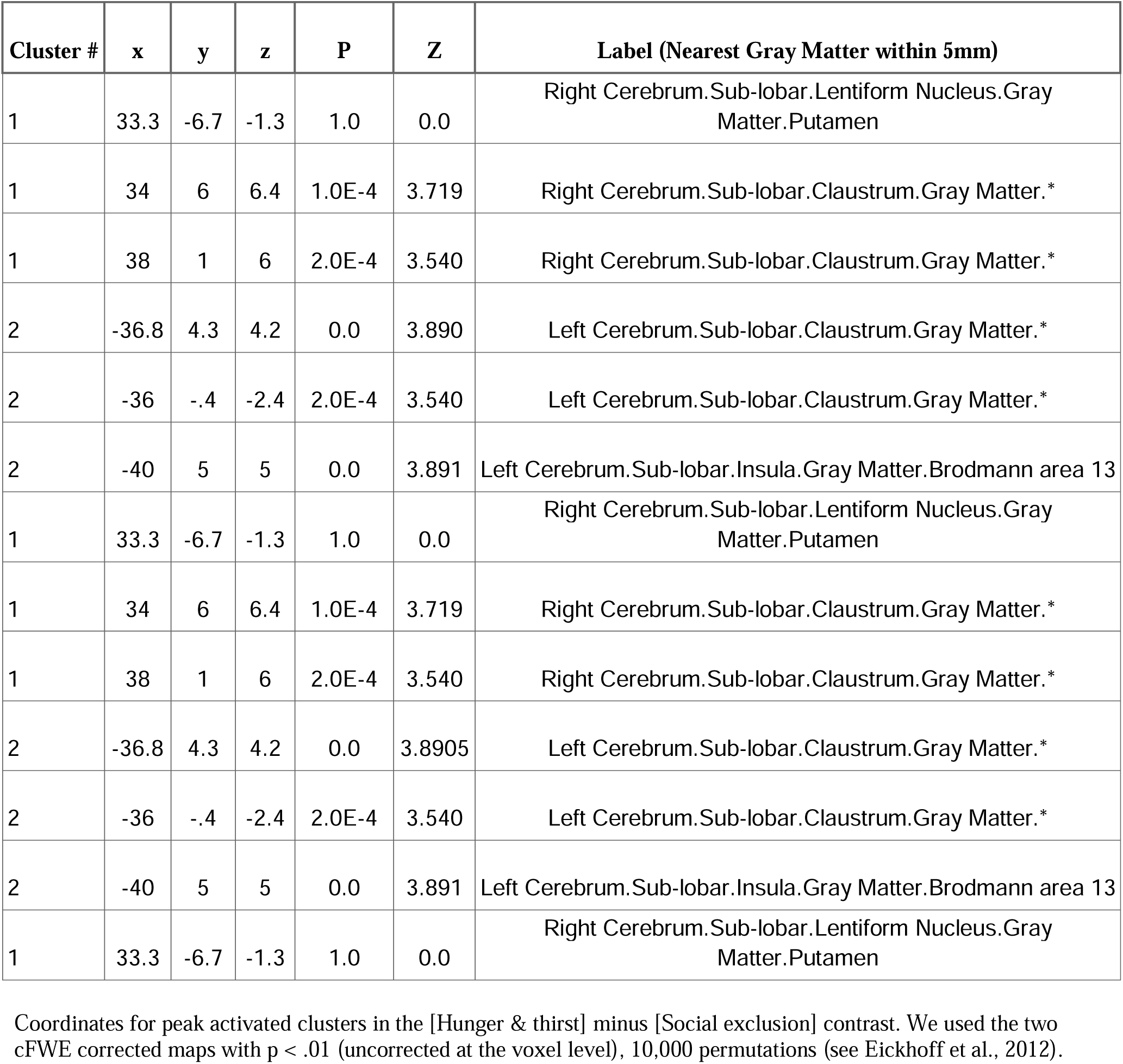
[Hunger & thirst] minus [Social deprivation].

**Table 8.** [Social Exclusion] Minus [Hunger & Thirst].

| Cluster # | x | y | z | P | Z | Label (Nearest Gray Matter within 5mm) |
| --- | --- | --- | --- | --- | --- | --- |
| 1 | 41 | -15.7 | 20.4 | 0.0 | 3.891 | Right Cerebrum.Sub-lobar.Insula.Gray Matter.Brodmann area 13 |

Coordinates for peak activated clusters in the [Social exclusion] minus [Hunger & thirst] contrast. We used the two cFWE corrected maps with $p < .01$ (uncorrected at the voxel level), 10,000 permutations (see Eickhoff et al., 2012).

**Figure 2.**
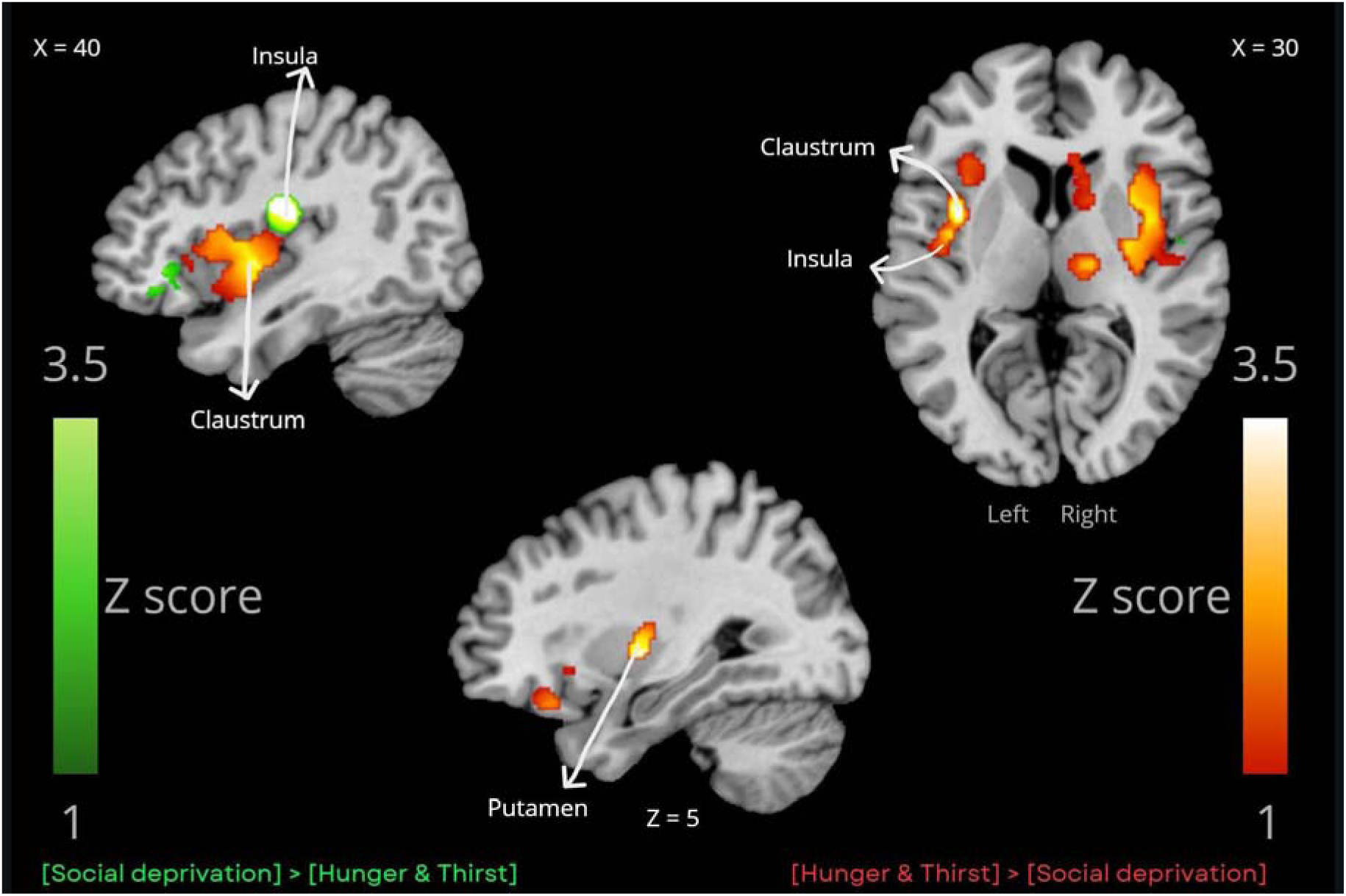
Contrasts Maps. In red, clustered thresholded maps for clusters of subtraction {[Physiological-Need] minus [Social-Need]}. In green, clustered thresholded maps for clusters of subtraction {[Social-Need] minus [Physiological-Need]}.

### Conjunction Meta-Analysis

The conjunction analysis of [Physiological-Need] AND [Social-Need] revealed overlapping consistent activation in the right posterior insula, right caudate head, and right ventral anterior cingulate cortex (ACC; BA24) (see Table 9 and Figure 3). The ACC cluster size was 8 mm³, which falls below the commonly used minimum threshold of 10 mm³.

**Table 9.** [Hunger & Thirst] and [Social Deprivation].

| Cluster # | x | y | z | ALE | Label (Nearest Gray Matter within 5mm) |
| --- | --- | --- | --- | --- | --- |
| 1 | 48 | -10 | 6 | 0.019 | Right Cerebrum.Sub-lobar.Insula.Gray Matter.Brodmann area 13 |
| 2 | 10 | 14 | -2 | 0.018 | Right Cerebrum.Sub-lobar.Caudate.Gray Matter.Caudate Head |
| 3 | 10 | 38 | -6 | 0.015 | Right Cerebrum.Limbic Lobe.Anterior Cingulate.Gray Matter.Brodmann area 24 |

Coordinates for the intersection between s in the [Hunger & thirst] condition AND [Social exclusion] condition. The conjunction was the intersection of the two cFWE thresholded maps.

**Figure 3.**
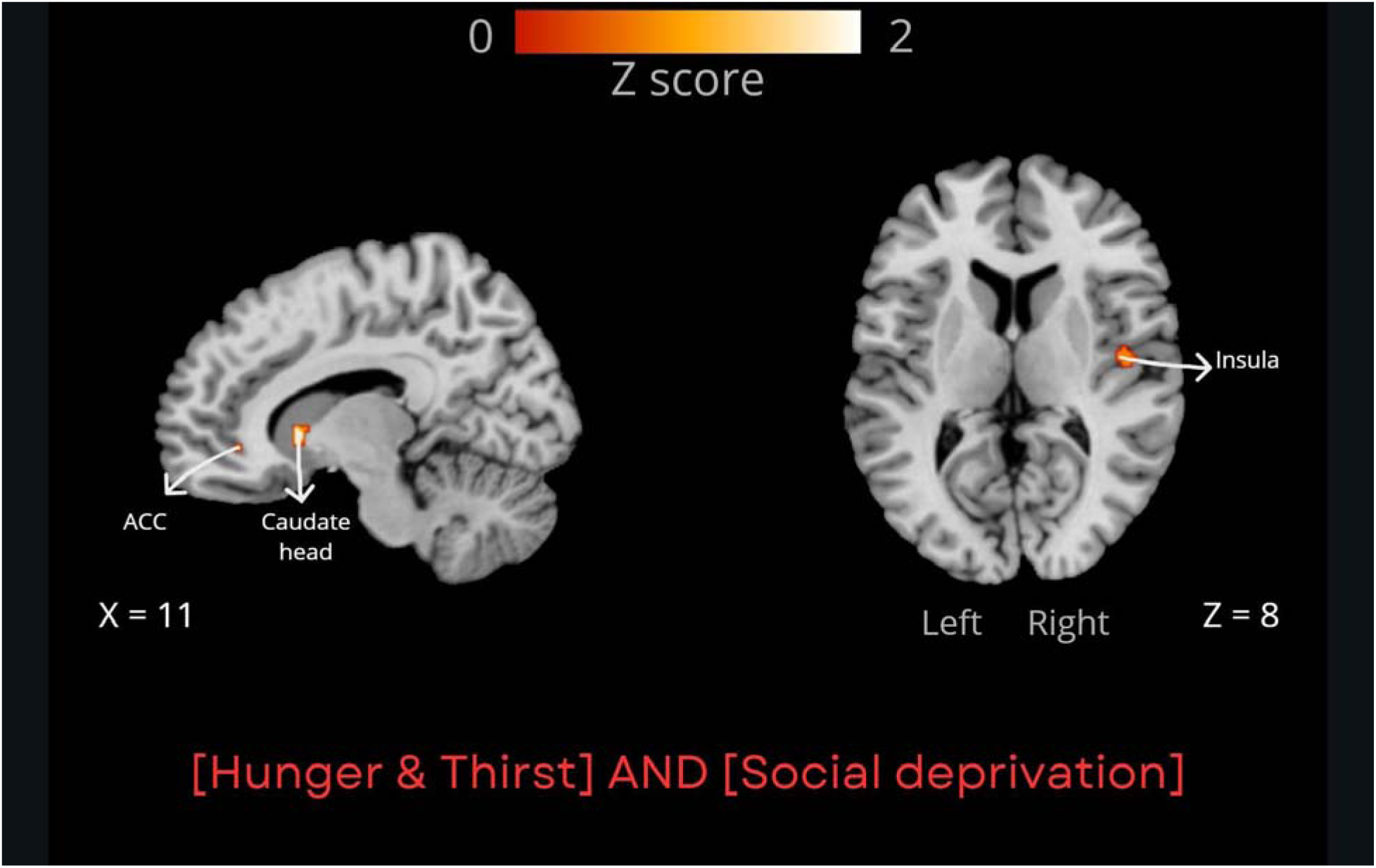
Conjunction maps. Clustered thresholded maps showing the intersection between activation patterns of [Physiological-Need] AND [Social-Need].

**Figure 4.**
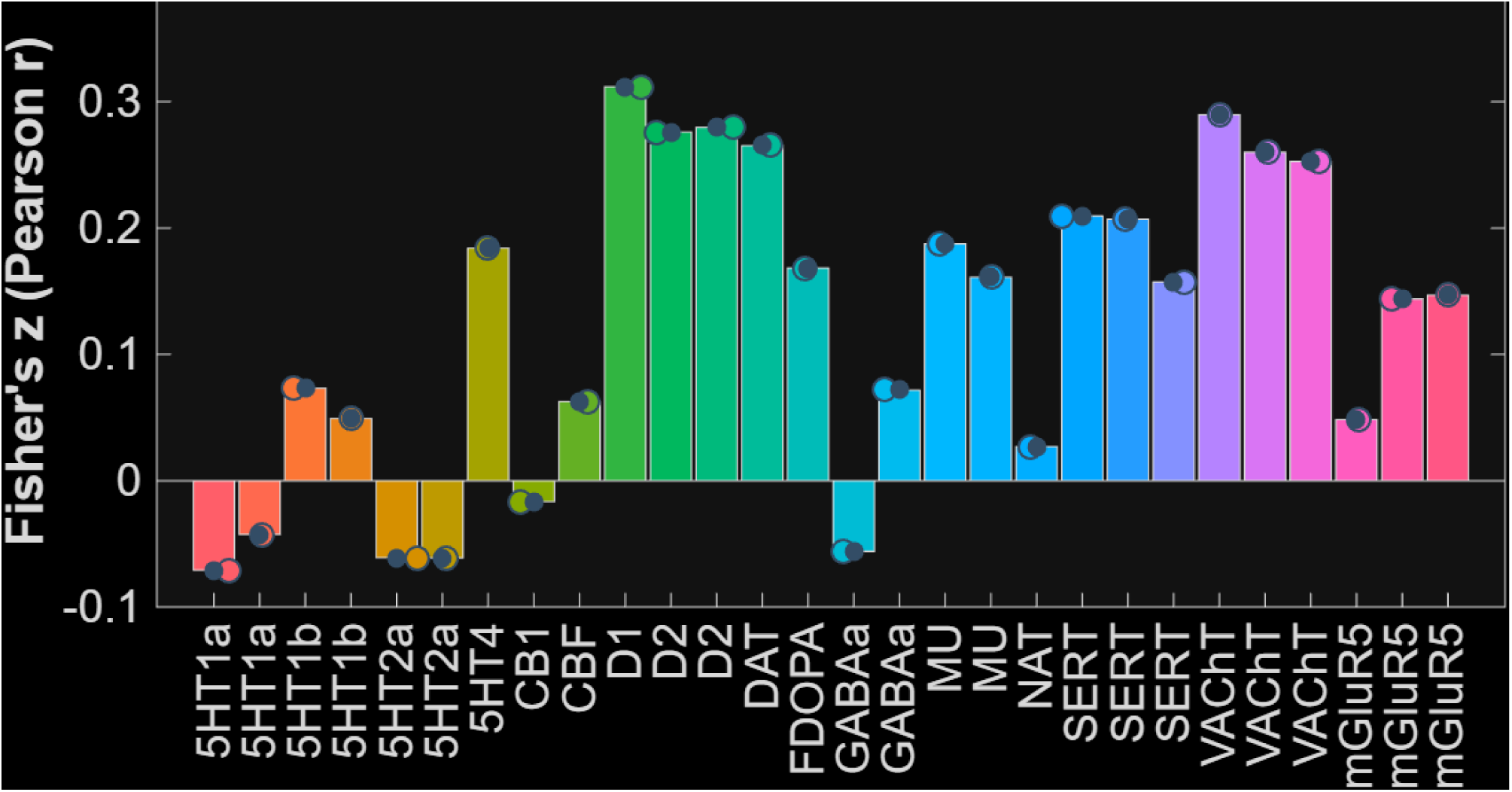
Spatial correlation between the conjunction map and serotonin receptors. Error bars showing the Fisher’s z Pearson correlation between the conjunction data on the y axis and the neurotransmitter map based on PET studies on the x axis. The colored dots represent data points and the black ones the mean of the bar which equals the to the single point.

### Spatial Correlation Between the Conjunction Map and Serotonin Neurotransmitter Receptors

Using JuSpace, we examined the spatial relationship between brain regions commonly activated by both physiological and social deprivation and whole-brain neurotransmitter maps. Given serotonin’s and dopamine’s role in food intake (van Galen et al., 2021; Baik, 2021), social exclusion (Preller et al., 2016; Chang et al., 2025), and aversive processing of salient stimuli (Dayan & Huys, 2009; Sizemore, 2020; Bromberg-Martin et al., 2010), we hypothesized an association with these shared activations.

While 5-HT4 had the highest correlation among serotonin receptors, it was not the highest overall (see Supplementary Material for full results). Positive correlations were also observed for D1 and D2 dopamine receptors, DAT (dopamine transporter), VAChT (acetylcholine transporter), SERT (serotonin transporter), 5-HT4 and 5-HT1B serotonin receptors, the μ-opioid receptor, and mGluR5 (metabotropic glutamate receptor). Negative correlations were found for 5-HT2A and 5-HT1A serotonin receptors, as well as CB1 endocannabinoid receptors. These findings suggest that dopamine, endogenous opioids, acetylcholine, and serotonin systems are spatially correlated with the intersection network.

## DISCUSSION

We investigated the shared (and distinct) brain activation patterns associated with physiological needs (hunger/thirst) and social exclusion. Using ALE meta-analysis, we compared consistent activation of stimulus perception under deprivation of the same stimulus: perceiving food/liquid during hunger/thirst and perceiving social interactions while being socially excluded. Separate meta-analyses revealed qualitative distinct activation patterns for physiological needs (insula, claustrum, putamen, ACC [Brodmann areas 24/25], caudate, parahippocampal gyrus, medial frontal gyrus, mammillary body, hippocampus) and social exclusion (insula, ACC [Brodmann areas 24/32], inferior frontal gyrus, OFC, anterior transverse temporal gyrus, caudate head). Contrast analyses showed greater activation for physiological deprivation within the putamen, claustrum, left posterior insula, OFC, and thalamus, mammillary body, while social deprivation elicited greater activation in the right posterior insula.

The aim of this study—to identify brain regions commonly engaged by hunger/thirst and social exclusion during the perception of need-relevant stimuli—was addressed using a conjunction analysis. This analysis revealed shared activation in the posterior insula, the caudate head and ventral ACC. Our results align with prior studies, which have reported activation of regions within the insula, striatum and ACC during perception of food during hunger (van der Lan et al., 2011; Goldstone et al., 2009; Bosulu et al., 2022), water during thirst (De Araujo et al., 2003), and social interaction during exclusion (Mwilambwe & Spreng, 2021). Subsequent JuSpace analyses showed that this shared network exhibited the strongest spatial correspondence with the 5-HT4 among serotonin receptors, whereas DA receptors were all highly associated with the shared network. Below, we discuss the implications of these findings with respect to the role of posterior insula, caudate head, and ventral ACC in stimulus processing under states of deprivation.

By analysing the conjunction of neural activity during social (exclusion) and physiological (hunger/thirst) deprivation, we offer novel insights into their shared neural substrates. Our findings reveal co-activation in the posterior insula, caudate head and ventral ACC—regions linked to aversive affect, motivation, and goal-directed behavior (Craig, 2003; Loonen & Ivanova, 2018; Balleine & O’Doherty, 2010). These findings support the idea that social exclusion shares some of the same neural systems that regulate interoception and homeostatic needs such as energy and fluid balance (Matthew & Tye, 2019; Craig, 2002). The posterior insula integrates emotional and homeostatic information, signaling the gap between actual and desired states (Craig, 2003; Gehrlach et al., 2019; Livneh et al., 2020; Barrett & Simmons, 2015). (2) and the ACC drives motivation, signaling the need for fulfillment to other brain regions, including the caudate (Craig, 2003; Weston, 2012; Peak et al., 2019); and the latter plays a role in goal directed behavior (Balleine and O’Doherty, 2010; Haber & Knutson, 2010; Hollon et al., 2014; ito and Doya, 2015; Schwabe & Wolf, 2010), based on the current need (van den Bercken & Cools, 1982).

Our findings align with recent literature in several important ways. Although social exclusion has often been linked to dorsal ACC activation, akin to physical pain (Eisenberger, 2012), we instead observed consistent ventral ACC activation, in line with a recent Cyberball meta-analysis (Mwilambwe & Spreng, 2021). Moreover, Tomova et al. (2020) reported that hunger-related food perception and post-isolation (but not exclusion) social interaction engaged dopaminergic SN/VTA regions associated with reward prediction and motivation. SN/VTA projects dopamine to the dorsal/ventral striatum, including to the caudate and the nucleus accumbens (Haber & Knutson, 2010). Indeed, those regions have been shown to be important for motivation and goal directed behavior. Thus, while our exclusion-focused conjunction analysis did not reveal SN/VTA activation, we did observe striatal engagement, a primary target of VTA/SN dopamine implicated in action and reward seeking. All this taken together might mean that social exclusion shares some deeper links with physiological needs.

Beyond identifying shared activation for physiological and social deprivation, we explored the relationship between these common regions (posterior insula, caudate head, and ventral ACC) and neurotransmitter receptor distribution, focusing on DA and 5-HT. While all DA receptors seemed to be similarly associated with the shared network, we found the strongest association between these regions and the 5-HT4 among serotonergic receptors. The presence of 5-HT4 is likely influenced by the presence of the caudate head in the shared network (Rebholz et al., 2018). In the mouse dorsal striatum, dopamine (DA) and serotonin (5-HT) act as complementary neuromodulatory systems via D1 and 5-HT4 receptors, respectively (Taylor et al., 2020). This organization may enable selective tuning of striatal neuron activity, supporting reinforcement, reward processing, and behavioral flexibility (Peters et al., 2021). Additionally, the medial PFC (BA32/25, within the ACC (Weston, 2012)) can influence activity in the serotonergic dorsal raphe (DRN) through a feedback loop (Rebholz et al., 2018; Peyron et al., 1997; Lucas et al., 2005; Price, 2007; Weston, 2012) that includes 5-HT4 as key component (Rebholz et al., 2018). This feedback loop is said to be crucial for affective motivation and internal model updating (Craig, 2003; Kolling et al., 2016; Petzschner et al., 2021). Thus, ventral ACC may, via 5-HT4, regulate DRN activity, ultimately influencing VTA/SN (Gervais & Rouillard, 2000). In short, our correlational findings and our interpretation suggest that perceiving stimuli one is deprived of engages brain regions that modulate dopamine and serotonin neuron activity.

## LIMITS

The main limit of our study is that for physiological needs we only examined hunger and thirst, while for social needs, we focused on exclusion, not isolation. Nevertheless, hunger and thirst share a common neural network (including the ACC and insula) similar to other homeostatic needs (McKinley et al., 2019; Craig, 2003); and while distinct, social exclusion (being outcast) and isolation (lack of contact) both represent a “lack of ties” (Huisman & van Tilburg, 2021). Exclusion, as used here, may represent short-term social deprivation (Cacioppo et al., 2002). Nevertheless, the Cyberball exclusion paradigm used here has been shown to induce need-like distress (Bernstein & Claypool, 2012), despite some debate about its effects (Gerber & Wheeler, 2009; Blackhart et al., 2009). Future research should explore a wider range of physiological and social deprivations. Finally, the limitations of reverse inference should be considered in interpreting our findings (Poldrack, 2006, 2011).

## CONCLUSION

Our goal was to study the neural substrates shared by social exclusion and physiological needs. Our meta-analysis reveals distinct and shared neural correlates for physiological (hunger/thirst) and social (exclusion) deprivation. While distinct networks exist, shared activation in the posterior insula, caudate head, and ACC suggests that brain mechanisms related to perception of social interaction during social exclusion have similarities to those related to perception of food/water when hungry/thirsty. Furthermore, these shared regions correlate significantly with dopamine and 5-HT4 receptors distribution. Overall, our findings suggest that perceiving deprivation-alleviating stimuli engages brain regions related to aversive states, goal-directed behavior, and potentially dopaminergic and serotonergic activity. This highlights a neural bridge between basic physiological drives and complex social needs, offering new insights into the architecture of human affective states.

## ACKNOWLEDGMENTS

The research was supported in part by NSERC Discovery Grant #RGPIN-2018-05698 and UdeM institutional funds.

## AUTHOR CONTRIBUTION

**Juvénal Bosulu**: Designed the study, performed the database search, performed data analysis, interpretation, and wrote the manuscript. **Yousra Mzireg :** Performed a database search, performed data analysis and revised the manuscript. **Sébastien Hétu**: Revised the manuscript and provided critical feedback. **Yi Luo**: Revised the manuscript and provided critical feedback. All authors contributed to and approved the final manuscript version.

## DATA AVAILABILITY STATEMENT

All Data are available upon request.

## REFERENCES

Achterberg, E. M., van Swieten, M. M., Driel, N. V., Trezza, V., & Vanderschuren, L. J. (2016). Dissociating the role of endocannabinoids in the pleasurable and motivational properties of social play behaviour in rats. Pharmacological Research, 110, 151–158. 10.1016/j.phrs.2016.04.031

Bach, P., Frischknecht, U., Bungert, M., Karl, D., Vollmert, C., Vollstädt-Klein, S., Lis, S., Kiefer, F., & Hermann, D. (2019). Effects of social exclusion and physical pain in chronic opioid maintenance treatment: fMRI correlates. European Neuropsychopharmacology, 29(2), 291–305. 10.1016/j.euroneuro.2018.11.1109

Baik, J. (2021). Dopaminergic control of the feeding circuit. Endocrinology and Metabolism, 36(2), 229–239. 10.3803/enm.2021.979

Balleine, B. W., & O’Doherty, J. P. (2010). Human and rodent homologies in action control: Corticostriatal determinants of goal-directed and habitual action. Neuropsychopharmacology, 35(1), 48–69. 10.1038/npp.2009.131

Barrett, L. F., & Simmons, W. K. (2015). Interoceptive predictions in the brain. Nature Reviews Neuroscience, 16(7), 419–429. 10.1038/nrn3950

Baumeister, R. F., & Leary, M. R. (1995). The need to belong: Desire for interpersonal attachments as a fundamental human motivation. Psychological Bulletin, 117(3), 497–529. 10.1037/0033-2909.117.3.497

Becker, C. A., Schmälzle, R., Flaisch, T., Renner, B., & Schupp, H. T. (2015). Thirst and the state-dependent representation of incentive stimulus value in human motive circuitry. Social Cognitive and Affective Neuroscience, 10(12), 1722–1729. 10.1093/scan/nsv063

Becker, C. A., Flaisch, T., Renner, B., & Schupp, H. T. (2017). From thirst to satiety: The anterior mid-cingulate cortex and right posterior insula indicate dynamic changes in incentive value. Frontiers in Human Neuroscience, 11, 234. 10.3389/fnhum.2017.00234

Bernstein, M. J., & Claypool, H. M. (2012). Not all social exclusions are created equal: Emotional distress following social exclusion is moderated by the exclusion paradigm. Social Influence, 7(2), 113–130. 10.1080/15534510.2012.664326

Bindra, D. (1974). A motivational view of learning, performance, and behavior modification. Psychological Review, 81(3), 199–213. 10.1037/h0036330

Blackhart, G. C., Nelson, B. C., Knowles, M. L., & Baumeister, R. F. (2009). Rejection elicits emotional reactions but neither causes immediate distress nor lowers self-esteem: A meta-analytic review of 192 studies on social exclusion. Personality and Social Psychology Review, 13(4), 269–309. 10.1177/1088868309346065

Bolling, D. Z., Pitskel, N. B., Deen, B., Crowley, M. J., McPartland, J. C., Mayes, L. C., & Pelphrey, K. A. (2011a). Dissociable brain mechanisms for processing social exclusion and rule violation. NeuroImage, 54(3), 2462–2471. 10.1016/j.neuroimage.2010.10.049

Bolling, D. Z., Pitskel, N. B., Deen, B., Crowley, M. J., Mayes, L. C., & Pelphrey, K. A. (2011b). Development of neural systems for processing social exclusion from childhood to adolescence. Developmental Science, 14(6), 1431–1444. 10.1111/j.1467-7687.2011.01087.x

Bolling, D. Z., Pitskel, N. B., Deen, B., Crowley, M. J., McPartland, J. C., Kaiser, M. D., Vander Wyk, B. C., Wu, J., Mayes, L. C., & Pelphrey, K. A. (2011c). Enhanced neural responses to rule violation in children with autism: A comparison to social exclusion. Developmental Cognitive Neuroscience, 1(3), 280–294. 10.1016/j.dcn.2011.02.002

Bolling, D. Z., Pelphrey, K. A., & Vander Wyk, B. C. (2015). Trait-level temporal lobe hypoactivation to social exclusion in unaffected siblings of children and adolescents with autism spectrum disorders. Developmental Cognitive Neuroscience, 13, 75–83. 10.1016/j.dcn.2015.04.007

Bosulu, J., Allaire, M., Tremblay-Grénier, L., Luo, Y., Eickhoff, S., & Hétu, S. (2022). “Wanting” versus “needing” related value: An fMRI meta-analysis. Brain and Behavior, 12(9), e32713. 10.1002/brb3.2713

Boureau, Y. L., & Dayan, P. (2011). Opponency revisited: Competition and cooperation between dopamine and serotonin. Neuropsychopharmacology, 36(1), 74–97. 10.1038/npp.2010.151

Bromberg-Martin, E. S., Matsumoto, M., & Hikosaka, O. (2010). Dopamine in motivational control: Rewarding, aversive, and alerting. Neuron, 68(5), 815–834. 10.1016/j.neuron.2010.11.022

Cacioppo, J. T., Ernst, J. M., Burleson, M. H., McClintock, M. K., Malarkey, W. B., Hawkley, L. C., … Berntson, G. G. (2000). Lonely traits and concomitant physiological processes: The MacArthur social neuroscience studies. International Journal of Psychophysiology, 35(2–3), 143–154. 10.1016/S0167-8760(99)00049-5

Cacioppo, J. T., Hawkley, L. C., Crawford, L. E., Ernst, J. M., Burleson, M. H., Kowalewski, R. B., … Berntson, G. G. (2002). Loneliness and health: Potential mechanisms. Psychosomatic Medicine, 64(3), 407–417. 10.1097/00006842-200205000-00005

Chang, H., Cheng, K., Hung, Y., & Hsu, K. (2025). Oxytocin signaling in the ventral tegmental area mediates social isolation-induced craving for social interaction. Journal of Biomedical Science, 32(1), 37. 10.1186/s12929-025-01130-0

Cheah, Y. S., Lee, S., Ashoor, G., Nathan, Y., Reed, L. J., Zelaya, F. O., … Amiel, S. A. (2014). Ageing diminishes the modulation of human brain responses to visual food cues by meal ingestion. International Journal of Obesity, 38(9), 1186–1192. 10.1038/ijo.2013.237

Chester, D. S., Lynam, D. R., Milich, R., & DeWall, C. N. (2018). Neural mechanisms of the rejection–aggression link. Social Cognitive and Affective Neuroscience, 13(5), 501–512. 10.1093/scan/nsy025

Cogoni, C., Carnaghi, A., & Silani, G. (2018). Reduced empathic responses for sexually objectified women: An fMRI investigation. Cortex, 99, 258–272. 10.1016/j.cortex.2017.11.020

Craig, A. D. (2002). How do you feel? Interoception: The sense of the physiological condition of the body. Nature Reviews Neuroscience, 3, 655–666. 10.1038/nrn894

Dayan, P., & Huys, Q. J. (2009). Serotonin in affective control. Annual Review of Neuroscience, 32(1), 95–126. 10.1146/annurev.neuro.051508.135607

De Araujo, I. E., Kringelbach, M. L., Rolls, E. T., & McGlone, F. (2003). Human cortical responses to water in the mouth, and the effects of thirst. Journal of Neurophysiology, 90(3), 1865–1876. 10.1152/jn.00297.2003

DeWall, C. N., Masten, C. L., Powell, C., Combs, D., Schurtz, D. R., & Eisenberger, N. I. (2012). Do neural responses to rejection depend on attachment style? An fMRI study. Social Cognitive and Affective Neuroscience, 7(2), 184–192. 10.1093/scan/nsq107

de Water, E., Mies, G. W., Ma, I., Mennes, M., Cillessen, A. H. N., & Scheres, A. (2017). Neural responses to social exclusion in adolescents: Effects of peer status. Cortex, 92, 32–43. 10.1016/j.cortex.2017.02.018

Dukart, J., Holiga, S., Rullmann, M., Lanzenberger, R., Hawkins, P. C. T., Mehta, M. A., … Eickhoff, S. B. (2021). JuSpace: A tool for spatial correlation analyses of magnetic resonance imaging data with nuclear imaging derived neurotransmitter maps. Human Brain Mapping, 42(3), 555–566. 10.1002/hbm.25244

Eklund, A., Nichols, T. E., & Knutsson, H. (2016). Cluster failure: Why fMRI inferences for spatial extent have inflated false-positive rates. Proceedings of the National Academy of Sciences of the United States of America, 113(28), 7900–7905. 10.1073/pnas.1602413113

Egan, G., Silk, T., Zamarripa, F., Williams, J., Federico, P., Cunnington, R., … Denton, D. A. (2003). Neural correlates of the emergence of consciousness of thirst. Proceedings of the National Academy of Sciences, 100(25), 15241–15246. 10.1073/pnas.2136650100

Eickhoff, S. B., Laird, A. R., Grefkes, C., Wang, L. E., Zilles, K., & Fox, P. T. (2009). Coordinate-based ALE meta-analysis of neuroimaging data: A random-effects approach based on empirical estimates of spatial uncertainty. Human Brain Mapping, 30(9), 2907–2926. 10.1002/hbm.20718

Eickhoff, S. B., Bzdok, D., Laird, A. R., Kurth, F., & Fox, P. T. (2012). Activation likelihood estimation meta analysis revisited. NeuroImage, 59(3), 2349–2361. 10.1016/j.neuroimage.2011.09.017

Eisenberger, N. I., Lieberman, M. D., & Williams, K. D. (2003). Does rejection hurt? An fMRI study of social exclusion. Science, 302(5643), 290–292. 10.1126/science.1089134

Eisenberger, N. I. (2012). The pain of social disconnection: Examining the shared neural underpinnings of physical and social pain. Nature Reviews Neuroscience, 13(6), 421–434. 10.1038/nrn3231

Falk, E. B., Cascio, C. N., O’Donnell, M. B., Carp, J., Tinney, F. J., Jr., Bingham, C. R., Shope, J. T., Ouimet, M. C., Pradhan, A. K., & Simons-Morton, B. G. (2014). Neural responses to exclusion predict susceptibility to social influence. Journal of Adolescent Health, *54*(5 Suppl), S22–S31. 10.1016/j.jadohealth.2013.12.035

Farrell, M. J., Bowala, T. K., Gavrilescu, M., Phillips, P. A., McKinley, M. J., McAllen, R. M., … Egan, G. F. (2011). Cortical activation and lamina terminalis functional connectivity during thirst and drinking in humans. American Journal of Physiology Regulatory, Integrative and Comparative Physiology, 301(3), R623–R631. 10.1152/ajpregu.00817.2010

Faurion, A., Cerf, B., Van De Moortele, P. F., Lobel, E., Mac Leod, P., & Le Bihan, D. (1999). Human taste cortical areas studied with functional magnetic resonance imaging: Evidence of functional lateralization related to handedness. Neuroscience Letters, 277(3), 189–192. 10.1016/s0304-3940(99)00881-2

Frank, S., Laharnar, N., Kullmann, S., Veit, R., Canova, C., Hegner, Y. L., … & Preissl, H. (2010). Processing of food pictures: Influence of hunger, gender and calorie content. Brain Research, 1350, 159–166. 10.1016/j.brainres.2010.04.030

Führer, D., Zysset, S., & Stumvoll, M. (2008). Brain activity in hunger and satiety: An exploratory visually stimulated fMRI study. Obesity, 16(5), 945–950. 10.1038/oby.2008.33

Gehrlach, D. A., Dolensek, N., Klein, A. S., Chowdhury, R. R., Matthys, A., Junghänel, M., Gaitanos, T. N., Podgornik, A., Black, T. D., Vaka, N. R., Conzelmann, K.-K., & Gogolla, N. (2020). Aversive state processing in the posterior insular cortex. Nature Neuroscience, 22(9), 1424–1437. 10.1038/s41593-019-0469-1

Gerber, J., & Wheeler, L. (2009). On being rejected: A meta analysis of experimental research on rejection. Perspectives on Psychological Science, 4(5), 468–488. 10.1111/j.1745-6924.2009.01158.x

Gervais, J., & Rouillard, C. (2000). Dorsal raphe stimulation differentially modulates dopaminergic neurons in the ventral tegmental area and substantia nigra. Synapse, 35(4), 281– 291. 10.1002/(SICI)1098-2396(20000315)35:4<281::AID-SYN6>3.0.CO;2-A

Goebel, B. L., & Brown, D. R. (1981). Age differences in motivation related to Maslow’s need hierarchy. Developmental Psychology, 17(6), 809–815. 10.1037/0012-1649.17.6.809

Goldstone, A. P., Prechtl de Hernandez, C. G., Beaver, J. D., Muhammed, K., Croese, C., Bell, G., … & Bell, J. D. (2009). Fasting biases brain reward systems towards high calorie foods. European Journal of Neuroscience, 30(8), 1625–1635. 10.1111/j.1460-9568.2009.06949.x

Gilman, J. M., Curran, M. T., Calderon, V., Schuster, R. M., & Evins, A. E. (2016). Altered neural processing to social exclusion in young adult marijuana users. Biological Psychiatry: Cognitive Neuroscience and Neuroimaging, 1(2), 152–159. 10.1016/j.bpsc.2015.11.002

Gonzalez, M. Z., Beckes, L., Chango, J., Allen, J. P., & Coan, J. A. (2015). Adolescent neighborhood quality predicts adult dACC response to social exclusion. Social Cognitive and Affective Neuroscience, 10(7), 921–928. 10.1093/scan/nsu137

Green, E., Jacobson, A., Haase, L., & Murphy, C. (2015). Neural correlates of taste and pleasantness evaluation in the metabolic syndrome. Brain Research, 1620, 57–71. 10.1016/j.brainres.2015.03.034

Gradin, V. B., Waiter, G., Kumar, P., Stickle, C., Milders, M., Matthews, K., … Steele, J. D. (2012). Abnormal neural responses to social exclusion in schizophrenia. PLoS ONE, 7(8), e42608. 10.1371/journal.pone.0042608

Haase, L., Cerf Ducastel, B., & Murphy, C. (2009). Cortical activation in response to pure taste stimuli during the physiological states of hunger and satiety. NeuroImage, 44(3), 1008–1021. 10.1016/j.neuroimage.2008.09.044

Haase, L., Green, E., & Murphy, C. (2011). Males and females show differential brain activation to taste when hungry and sated in gustatory and reward areas. Appetite, 57(2), 421–434. 10.1016/j.appet.2011.06.009

Haber, S. N., & Knutson, B. (2010). The reward circuit: linking primate anatomy and human imaging. Neuropsychopharmacology, 35(1), 4–26. 10.1038/npp.2009.129

Hamid, A. A., Pettibone, J. R., Mabrouk, O. S., Hetrick, V. L., Schmidt, R., Vander Weele, C. M., … & Berke, J. D. (2016). Mesolimbic dopamine signals the value of work. Nature Neuroscience, 19(1), 117–126. 10.1038/nn.4173

Harding, I. H., Andrews, Z. B., Mata, F., Orlandea, S., Martinez Zalacain, I., Soriano Mas, C., … Verdejo Garcia, A. (2018). Brain substrates of unhealthy versus healthy food choices: Influence of homeostatic status and body mass index. International Journal of Obesity, 42(3), 448–456. 10.1038/ijo.2017.237

He, Q., Huang, X., Zhang, S., Turel, O., Ma, L., & Bechara, A. (2019). Dynamic causal modeling of insular, striatal, and prefrontal cortex activities during a food specific Go/NoGo task. Biological Psychiatry: Cognitive Neuroscience and Neuroimaging, 4(12), 1080–1089. 10.1016/j.bpsc.2018.12.005

Hollon, N. G., Arnold, M. M., Gan, J. O., Walton, M. E., & Phillips, P. E. (2014). Dopamine-associated cached values are not sufficient as the basis for action selection. Proceedings of the National Academy of Sciences, 111(51), 18357–18362. 10.1073/pnas.1419770111

Holsen, L. M., Zarcone, J. R., Thompson, T. I., Brooks, W. M., Anderson, M. F., Ahluwalia, J. S., … Savage, C. R. (2005). Neural mechanisms underlying food motivation in children and adolescents. NeuroImage, 27(3), 669–676. 10.1016/j.neuroimage.2005.04.043

Holsen, L. M., Zarcone, J. R., Brooks, W. M., Butler, M. G., Thompson, T. I., Ahluwalia, J. S., … Savage, C. R. (2006). Neural mechanisms underlying hyperphagia in Prader Willi syndrome. Obesity, 14(6), 1028–1037. 10.1038/oby.2006.118

Hofstede, G. (1984). Culture’s consequences: International differences in work-related values (Vol. 5). Sage.

Huisman, M., & van Tilburg, T. G. (2021). Social exclusion and social isolation in later life. In Handbook of aging and the social sciences (pp. 99–114). Academic Press. 10.1016/B978-0-12-815970-5.00007-3

Hull, C. L. (1943). Principles of behavior: An introduction to behavior theory. Appleton-Century.

Ito, M., & Doya, K. (2015). Distinct neural representation in the dorsolateral, dorsomedial, and ventral parts of the striatum during fixed and free choice tasks. Journal of Neuroscience, 35(8), 3499–3514. 10.1523/JNEUROSCI.1962-14.2015

Jacobson, A., Green, E., & Murphy, C. (2010). Age related functional changes in gustatory and reward processing regions: An fMRI study. NeuroImage, 53(2), 602–610. 10.1016/j.neuroimage.2010.05.012

Jacobson, A., Green, E., Haase, L., Szajer, J., & Murphy, C. (2019). Differential effects of BMI on brain response to odor in olfactory, reward and memory regions: Evidence from fMRI. Nutrients, 11(4), 926. 10.3390/nu11040926

Jiang, T., Soussignan, R., Schaal, B., & Royet, J. P. (2014). Reward for food odors: an fMRI study of liking and wanting as a function of metabolic state and BMI. Social Cognitive and Affective Neuroscience, 10(4), 561–568. 10.1093/scan/nsu086

Kerem, L., Holsen, L., Fazeli, P., Bredella, M. A., Mancuso, C., Resulaj, M., Holmes, T. M., Klibanski, A., & Lawson, E. A. (2021). Modulation of neural fMRI responses to visual food cues by overeating and fasting interventions: A preliminary study. Physiological Reports, 8(24), e14639. 10.14814/phy2.14639

Knutson, B., & Cooper, J. C. (2005). Functional magnetic resonance imaging of reward prediction. Current Opinion in Neurology, 18(4), 411–417. 10.1097/01.wco.0000173463.24758.f6

Kolling, N., Wittmann, M. K., Behrens, T. E. J., Boorman, E. D., Mars, R. B., & Rushworth, M. F. S. (2016). Anterior cingulate cortex and the value of the environment, search, persistence, and model updating. Nature Neuroscience, 19(10), 1280. 10.1038/nn.4382

LaBar, K. S., Gitelman, D. R., Parrish, T. B., Kim, Y. H., Nobre, A. C., & Mesulam, M. (2001). Hunger selectively modulates corticolimbic activation to food stimuli in humans. Behavioral Neuroscience, 115(2), 493–503. 10.1037/0735-7044.115.2.493

Lancaster, J. L., Tordesillas Gutiérrez, D., Martinez, M., Salinas, F., Evans, A., Zilles, K., … Fox, P. T. (2007). Bias between MNI and Talairach coordinates analyzed using the ICBM 152 brain template. Human Brain Mapping, 28(11), 1194–1205. 10.1002/hbm.20345

Le, T. M., Zhornitsky, S., Wang, W., & Li, C. S. R. (2020). Perceived burdensomeness and neural responses to ostracism in the Cyberball task. Journal of Psychiatric Research, 130, 1–8. 10.1016/j.jpsychires.2020.06.015

Liu, Z., Lin, R., & Luo, M. (2020). Reward contributions to serotonergic functions. Annual Review of Neuroscience, 43(1), 141–162. 10.1146/annurev-neuro-093019-112252

Livneh, Y., Sugden, A. U., Madara, J. C., Essner, R. A., Flores, V. I., Sugden, L. A., … Andermann, M. L. (2020). Estimation of current and future physiological states in insular cortex. Neuron, 105(6), 1094–1111. 10.1016/j.neuron.2019.12.027

Loonen, A. J., & Ivanova, S. A. (2018). Circuits regulating pleasure and happiness: Evolution and role in mental disorders. Acta Neuropsychiatrica, 30(1), 29–42. 10.1017/neu.2017.8

Lucas, G., Compan, V., Charnay, Y., Neve, R. L., Nestler, E. J., Bockaert, J., … Debonnel, G. (2005). Frontocortical 5 HT4 receptors exert positive feedback on serotonergic activity: Viral transfections, subacute and chronic treatments with 5 HT4 agonists. Biological Psychiatry, 57(8), 918–925. 10.1016/j.biopsych.2004.12.023

Luo, M., Li, Y., & Zhong, W. (2016). Do dorsal raphe 5 HT neurons encode “beneficialness”? Neurobiology of Learning and Memory, 135, 40–49. 10.1016/j.nlm.2016.08.008

MacGregor, D. (1960). The human side of enterprise (Vol. 21, No. 166.1960). McGraw Hill: New York.

Maner, J. K., DeWall, C. N., Baumeister, R. F., & Schaller, M. (2007). Does social exclusion motivate interpersonal reconnection? Resolving the “porcupine problem.” Journal of Personality and Social Psychology, 92(1), 42–55. 10.1037/0022-3514.92.1.42

Martens, M. J., Born, J. M., Lemmens, S. G., Karhunen, L., Heinecke, A., Goebel, R., … & Westerterp-Plantenga, M. S. (2013). Increased sensitivity to food cues in the fasted state and decreased inhibitory control in the satiated state in the overweight. The American Journal of Clinical Nutrition, 97(3), 471–479. 10.3945/ajcn.112.044024

Maslow, A. H. (1943). A theory of human motivation. Psychological Review, 50(4), 370–396. 10.1037/h0054346

Masten, C. L., Eisenberger, N. I., Borofsky, L. A., Pfeifer, J. H., McNealy, K., Mazziotta, J. C., & Dapretto, M. (2009). Neural correlates of social exclusion during adolescence: Understanding the distress of peer rejection. Social Cognitive and Affective Neuroscience, 4(2), 143–157. 10.1093/scan/nsp007

McKinley, M. J., Denton, D. A., Ryan, P. J., Yao, S. T., Stefanidis, A., & Oldfield, B. J. (2019). From sensory circumventricular organs to cerebral cortex: Neural pathways controlling thirst and hunger. Journal of Neuroendocrinology, 31(3), e12689. 10.1111/jne.12689

Montague, P. R., Dayan, P., & Sejnowski, T. J. (1996). A framework for mesencephalic dopamine systems based on predictive Hebbian learning. Journal of Neuroscience, 16(5), 1936–1947. 10.1523/JNEUROSCI.16-05-01936.1996

Moor, B. G., Güroğlu, B., de Macks, Z. A. O., Rombouts, S. A. R. B., Van der Molen, M. W., & Crone, E. A. (2012). Social exclusion and punishment of excluders: Neural correlates and developmental trajectories. NeuroImage, 59(1), 708–717. 10.1016/j.neuroimage.2011.07.028

Mwilambwe Tshilobo, L., & Spreng, R. N. (2021). Social exclusion reliably engages the default network: A meta-analysis of Cyberball. NeuroImage, 227, 117666. 10.1016/j.neuroimage.2020.117666

Nishiyama, Y., Okamoto, Y., Kunisato, Y., Okada, G., Yoshimura, S., Kanai, Y., … Yamawaki, S. (2015). fMRI study of social anxiety during social ostracism with and without emotional support. PLOS ONE, 10(5), e0127426. 10.1371/journal.pone.0127426

Novembre, G., Zanon, M., & Silani, G. (2015). Empathy for social exclusion involves the sensory-discriminative component of pain: A within-subject fMRI study. Social Cognitive and Affective Neuroscience, 10(2), 153–164. 10.1093/scan/nsu038

Peak, J., Hart, G., & Balleine, B. W. (2019). From learning to action: The integration of dorsal striatal input and output pathways in instrumental conditioning. European Journal of Neuroscience, 49(5), 658–671. 10.1111/ejn.13964

Peyron, C., Petit, J. M., Rampon, C., Jouvet, M., & Luppi, P. H. (1997). Forebrain afferents to the rat dorsal raphe nucleus demonstrated by retrograde and anterograde tracing methods. Neuroscience, 82(2), 443–468. 10.1016/S0306-4522(97)00268-6

Petzschner, F. H., Garfinkel, S. N., Paulus, M. P., Koch, C., & Khalsa, S. S. (2021). Computational models of interoception and body regulation. Trends in Neurosciences, 44(1), 63– 76. 10.1016/j.tins.2020.09.012

Poldrack, R. A. (2006). Can cognitive processes be inferred from neuroimaging data? Trends in Cognitive Sciences, 10(2), 59–63. 10.1016/j.tics.2005.12.004

Poldrack, R. A. (2011). Inferring mental states from neuroimaging data: From reverse inference to large-scale decoding. Neuron, 72(5), 692–697. 10.1016/j.neuron.2011.11.001

Preller, K. H., Pokorny, T., Hock, A., Kraehenmann, R., Stämpfli, P., Seifritz, E., … & Vollenweider, F. X. (2016). Effects of serotonin 2A/1A receptor stimulation on social exclusion processing. Proceedings of the National Academy of Sciences, 113(18), 5119–5124. 10.1073/pnas.1524187113

Price, J. L. (2007). Definition of the orbital cortex in relation to specific connections with limbic and visceral structures and other cortical regions. Annals of the New York Academy of Sciences, 1121(1), 54–71. 10.1196/annals.1401.008

Puetz, V. B., Kohn, N., Dahmen, B., Zvyagintsev, M., Schüppen, A., Schultz, R. T., … Konrad, K. (2014). Neural response to social rejection in children with early separation experiences. Journal of the American Academy of Child & Adolescent Psychiatry, 53(12), 1328–1337. 10.1016/j.jaac.2014.09.004

Radke, S., Seidel, E. M., Boubela, R. N., Thaler, H., Metzler, H., Kryspin-Exner, I., … Derntl, B. (2018). Immediate and delayed neuroendocrine responses to social exclusion in males and females. Psychoneuroendocrinology, 93, 56–64. 10.1016/j.psyneuen.2018.04.005

Rebholz, H., Friedman, E., & Castello, J. (2018). Alterations of expression of the serotonin 5-HT4 receptor in brain disorders. International Journal of Molecular Sciences, 19(11), 3581. 10.3390/ijms19113581

Reeve, J., & Lee, W. (2018). A neuroscientific perspective on basic psychological needs. Journal of Personality, 87(1), 102–114. 10.1111/jopy.12390

Rice, M. E., Patel, J. C., & Cragg, S. J. (2011). Dopamine release in the basal ganglia. Neuroscience, 198, 112–137. 10.1016/j.neuroscience.2011.08.066

Saker, P., Farrell, M. J., Adib, F. R., Egan, G. F., McKinley, M. J., & Denton, D. A. (2014). Regional brain responses associated with drinking water during thirst and after its satiation. Proceedings of the National Academy of Sciences, 111(14), 5379–5384. 10.1073/pnas.1403382111

Saker, P., Farrell, M. J., Egan, G. F., McKinley, M. J., & Denton, D. A. (2018). Influence of anterior midcingulate cortex on drinking behavior during thirst and following satiation. Proceedings of the National Academy of Sciences of the United States of America, 115(4), 786–791. 10.1073/pnas.1717646115

Schultz, W. (1998). Predictive reward signal of dopamine neurons. Journal of Neurophysiology, 80(1), 1–27. 10.1152/jn.1998.80.1.1

Schultz, W., Dayan, P., & Montague, P. R. (1997). A neural substrate of prediction and reward. Science, 275(5306), 1593–1599. 10.1126/science.275.5306.1593

Schwabe, L., & Wolf, O. T. (2010). Learning under stress impairs memory formation. Neurobiology of Learning and Memory, 93(2), 183–188. 10.1016/j.nlm.2009.09.009

Siep, N., Roefs, A., Roebroeck, A., Havermans, R., Bonte, M. L., & Jansen, A. (2009). Hunger is the best spice: An fMRI study of the effects of attention, hunger, and calorie content on food reward processing in the amygdala and orbitofrontal cortex. Behavioural Brain Research, 198(1), 149–158. 10.1016/j.bbr.2008.10.035

Sizemore, T. R., Hurley, L. M., & Dacks, A. M. (2020). Serotonergic modulation across sensory modalities. Journal of Neurophysiology, 123(6), 2406–2425. 10.1152/jn.00034.2020

Tay, L., & Diener, E. (2011). Needs and subjective well-being around the world. Journal of Personality and Social Psychology, 101(2), 354–365. 10.1037/a0023779

Taylor, A. M., Ritchie, S. J., Madden, C., & Deary, I. J. (2020). Associations between Brief Resilience Scale scores and ageing-related domains in the Lothian Birth Cohort 1936. Psychology and Aging, 35(3), 329–344. 10.1037/pag0000419

Toates, F. (1994). Comparing motivational systems—An incentive motivation perspective. In C. R. Legg & D. A. Booth (Eds.), Appetite: Neural and behavioural bases (pp. 305–327). Oxford University Press. 10.1093/acprof:oso/9780198547877.003.0013

Tomova, L., Wang, K. L., Thompson, T., Matthews, G. A., Takahashi, A., Tye, K. M., & Saxe, R. (2020). Acute social isolation evokes midbrain craving responses similar to hunger. Nature Neuroscience, 23(12), 1597–1605. 10.1038/s41593-020-00742-z

Van den Bercken, J. H. L., & Cools, A. R. (1982). Evidence for a role of the caudate nucleus in the sequential organization of behaviour. Behavioural Brain Research, 4(4), 319–337. 10.1016/0166-4328(82)90058-4

van der Laan, L. N., De Ridder, D. T., Viergever, M. A., & Smeets, P. A. (2011). The first taste is always with the eyes: A meta-analysis on the neural correlates of processing visual food cues. NeuroImage, 55(1), 296–303. 10.1016/j.neuroimage.2010.11.055

van der Meulen, M., Steinbeis, N., Achterberg, M., van IJzendoorn, M. H., & Crone, E. A. (2018). Heritability of neural reactions to social exclusion and prosocial compensation in middle childhood. Developmental Cognitive Neuroscience, 34, 42–52. 10.1016/j.dcn.2018.05.010

van Galen, K. A., Ter Horst, K. W., & Serlie, M. J. (2021). Serotonin, food intake, and obesity. Obesity Reviews, 22(7), e13210. 10.1111/obr.13210

Vijayakumar, N., Cheng, T. W., & Pfeifer, J. H. (2017). Neural correlates of social exclusion across ages: A coordinate-based meta-analysis of functional MRI studies. NeuroImage, 153, 359–368. 10.1016/j.neuroimage.2017.02.050

Wagels, L., Bergs, R., Clemens, B., Bauchmüller, M., Gur, R. C., Schneider, F., … Kohn, N. (2017). Contextual exclusion processing: An fMRI study of rejection in a performance related context. Brain Imaging and Behavior, 11(3), 874–886. 10.1007/s11682-016-9561-2

Weston, C. S. (2012). Another major function of the anterior cingulate cortex: The representation of requirements. Neuroscience & Biobehavioral Reviews, 36(1), 90–110. 10.1016/j.neubiorev.2011.04.014

Will, G. J., van Lier, P. A., Crone, E. A., & Güroğlu, B. (2016). Chronic childhood peer rejection is associated with heightened neural responses to social exclusion during adolescence. Journal of Abnormal Child Psychology, 44(1), 43–55. 10.1007/s10802-015-9983-0

Williams, K. D. (2007). Ostracism. Annual Review of Psychology, 58, 425–452. 10.1146/annurev.psych.58.110405.085641

Williams, K. D., Cheung, C. K., & Choi, W. (2000). Cyberostracism: Effects of being ignored over the Internet. Journal of Personality and Social Psychology, 79(5), 748–762. 10.1037/0022-3514.79.5.748

Woo, C. W., Krishnan, A., & Wager, T. D. (2014). Cluster-extent based thresholding in fMRI analyses: pitfalls and recommendations. NeuroImage, 91, 412–419. 10.1016/j.neuroimage.2013.12.058

Wudarczyk, O. A., Kohn, N., Bergs, R., Gur, R. E., Turetsky, B. I., Schneider, F., & Habel, U. (2015). Chemosensory anxiety cues moderate the experience of social exclusion–an fMRI investigation with Cyberball. Frontiers in Psychology, 6, 1475. 10.3389/fpsyg.2015.01475

